# Spatial host-microbiome sequencing

**DOI:** 10.1101/2022.07.18.500470

**Authors:** Britta Lötstedt, Martin Stražar, Ramnik Xavier, Aviv Regev, Sanja Vickovic

## Abstract

Mucosal and barrier tissues such as the gut, lung or skin, are composed of a complex network of cells and microbes forming a tight niche that prevents pathogen colonization and supports host-microbiome symbiosis. Characterizing these networks at high molecular and cellular resolution is crucial for our understanding of homeostasis and disease. Spatial transcriptomics has emerged as a key technology to positionally profile RNAs at high resolution in tissues. Here, we present spatial host-microbiome sequencing, an all-sequencing based approach that captures tissue histology, polyadenylated RNAs and bacterial 16S sequences directly from tissues on spatially barcoded glass surfaces. We apply our approach to the mouse gut as a model system, use a novel deep learning approach for data mapping and detect spatial niches impacted by microbial biogeography. Spatial host-microbiome sequencing should enhance study of native host-microbe interactions in health and disease.

## INTRODUCTION

Mucosal and barrier tissues are ecosystems of multiple host cell types and a complex microbiome, and vary in space and time. Antigen recognition and innate immune responses[1] in the host, and molecular mechanisms derived from the microbiome, together prevent pathogen colonization and support the establishment of the host-microbiome spatial niche, and host-microbial symbiosis[2]. Conversely, spatial dysregulation[3,4] in diseases such as IBD can lead to dysfunction of the gut barrier[5,6].

Characterizing and understanding the host-microbiome spatial niche requires detailed measurement of the identity and molecular characteristics of host cells and microbiome species and their inter-relations in a spatial context. On the microbiome side, spatial metagenomics methods[7] are emerging to map bacteria by either imaging[8,9] or metagenomic plot sampling[10]. However, such studies focused on smaller regions, such as inter-fold, mucosal or lumen regions in the gut, and typically used broad taxonomy assignments, at best, reaching family level at best[10,11], with few reports at the level of specific genera or species[9,12–14]. Metagenomic plot-sampling, so far the only approach for spatial bacterial sequencing *in situ[10],* does not currently profile host gene expression. On the host side, single cell genomics, including single cell RNA-seq (scRNA-seq) has been instrumental to characterize the cellular composition of tissues, for either the host[15], and more recently, the resident microbes[16], but without spatial information. Spatial transcriptomics methods, either imaging- or sequencing-based, enable cell type mapping *in situ[17–20]*,^[21,22]^, but have not yet been applied to simultaneously profile both host and microbiome in a spatial context.

Here, we bridge this gap, by developing spatial host-microbiome sequencing (SHM-seq, **Fig1**), a robust all-sequencing based technology that leverages previous advancements in spatial transcriptomics[23,24], and provides histology, spatial RNA-seq and spatial 16S sequencing using readily available instrumentation to profile the host’s expression responses in relation to microbial biogeography. We apply it in the model system of the mouse colon, showing a roadmap for interrogating spatial gene expression programs in correlation to bacterial presence.

**Fig1.**
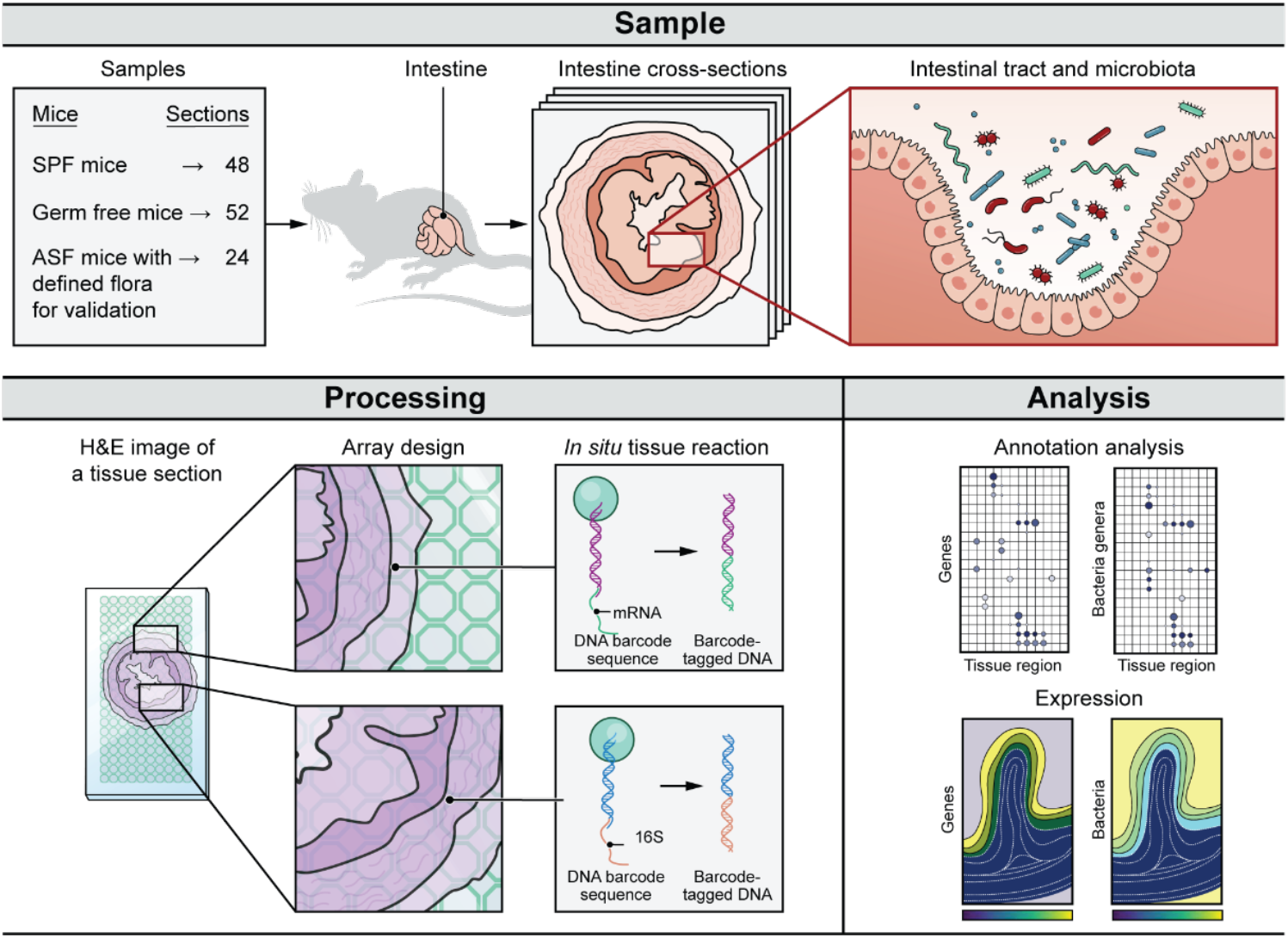
SHM-Seq: Spatial host-microbiome sequencing. Top: Three different mouse conditions used in the study analyzing cross sections in the mouse gut. Bottom left: Tissue sections from mouse colons were placed on a barcoded glass array, with a barcoded surface adapted for simultaneous capture of polyadenylated host transcripts and 16S bacterial rRNA. Tissue sections were imaged, cells permeabilized and cDNA synthesized on the array surface before library preparation and sequencing. Bottom right: Data analysis identifies regional gene programs, their cell-type constituents, association with mouse condition, and regional association with specific commensal bacteria.

## RESULTS

### Spatial host-microbiome sequencing

We developed SHM-seq by adapting Spatial Transcriptomics[23], where mRNA is captured by probes on a glass slide followed by profiling, to enable simultaneous capture of polyadenylated (host) transcripts and hypervariable regions of the 16S ribosomal RNA (rRNA) (**Methods, Fig1**). Specifically, we produced solid phase Spatial Transcriptomics slides covered with uniquely barcoded and spatially addressable capture probes[23–25], with ~1,000 distinct addressable DNA features (*i.e.* spatial spots) deposited and covalently linked to a glass substrate. We enzymatically modified these DNA features on the glass array to enable simultaneous capture of polyadenylated transcripts and hypervariable regions of the 16S rRNA[26] (**Methods, Fig1**). Next, we placed frozen tissue sections on the optimized glass surface, stained them with Hematoxylin and Eosin (H&E) and imaged the tissue histology by bright field microscopy. Finally, after imaging, we permeabilized the cells, allowing capture of host polyadenylated transcripts and bacterial 16S sequences on the array. The result was direct spatial DNA barcoding of host transcripts and bacterial species, which were sequenced by Illumina sequencing[24].

To test SHM-seq, we applied it to profile intestinal cross sections from the mouse colon (**Fig1**), of C57BL/6 mice grown under typical conditions (specific pathogen free; SPF), or as germ-free (GF) or altered schaedler flora (ASF) mice (**Methods**). Germ-free mice provide a negative control, whereas ASF mice provide a clear target for validation of the capture of expected bacterial species as they contain only a defined floral community and regular C57BL/6 SPF mice represent a complex case study with unaltered gut flora. In total, we applied SHM-seq to 124 tissue sections and collected data from 15,745 spatial spots across the three conditions.

### A deep learning approach for taxonomy classification of microbiome data

While Spatial Transcriptomics data has mature processing pipelines[27] (**Methods**), we devised a novel taxonomy assignment pipeline to process the spatial microbiome data. First, we created a customized bacterial reference (**Methods**) based on shotgun metagenomic sequencing data, comprising 65 species from 39 genera (**Fig2a**). Next, we devised an approach for taxonomy assignment, where spatially captured sequences were first classified using Kraken2[28] (**Methods**), and those without a taxonomic classification at the genus level were then processed by a novel deep learning approach (**Methods**). Our deep learning model (**Supplementary Fig1a**) is based on convolutional and recurrent neural networks, which process a read from both directions, seek local sequence patterns and their distant interactions, and are trained to predict the most likely genus assignment (**Methods**). We assessed its performance using simulated bacterial reads with attached taxa labels, mimicking data otherwise obtained with SHM-seq (**Methods**).

**Fig2.**
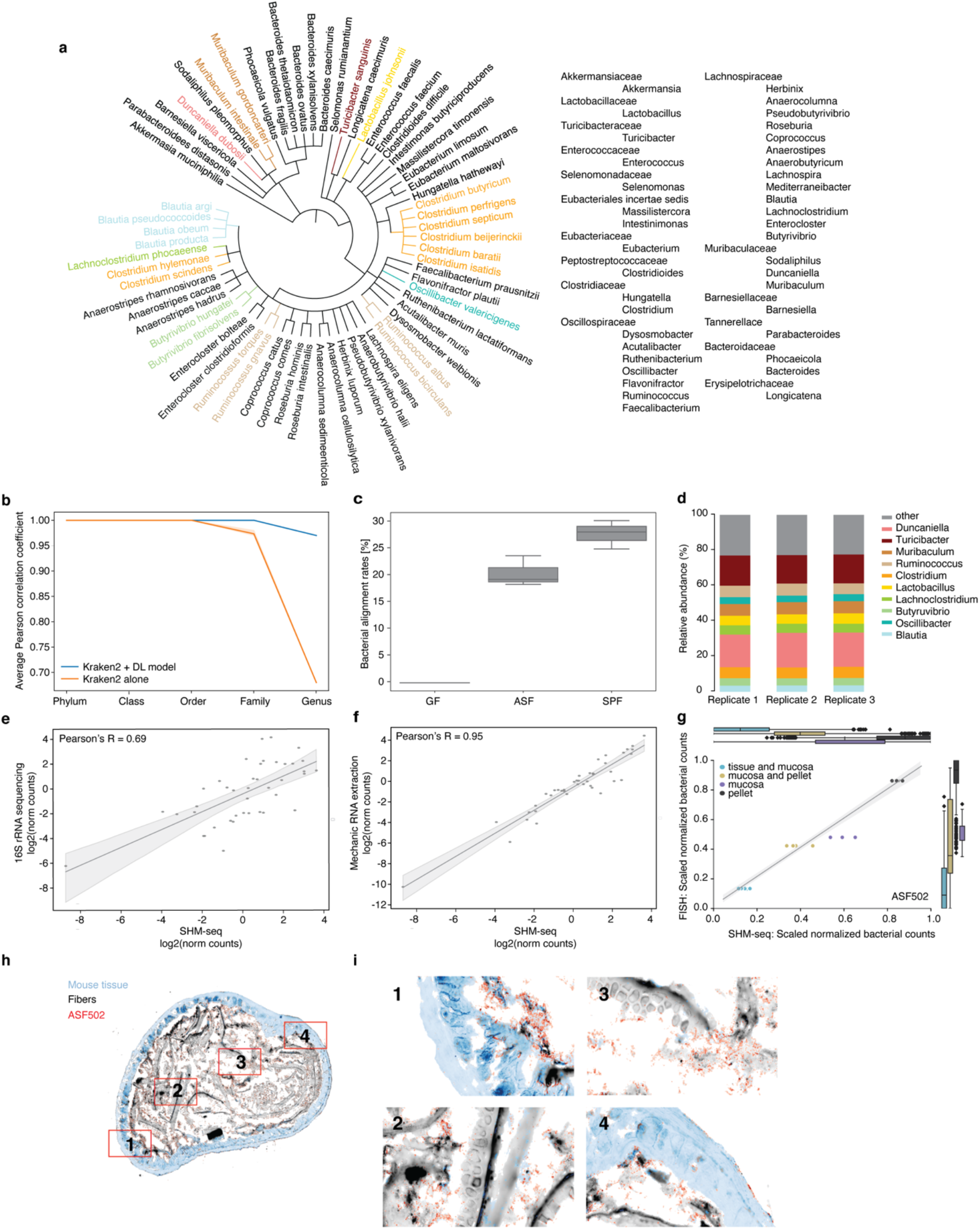
SHM-seq accurately captures bacterial representation and abundances in SPF and ASF mice. **(a)** Bacterial reference of the mouse gut microbiome. Phylogenetic tree based on metagenomic sequencing of colonic content from SPF mice, representing the 65 species in the mouse gut bacteria reference and colored to highlight taxonomic families and genera. **(b)** Enhanced annotation performance of the deep learning model. Average Pearson correlation coefficient (y axis) between true and predicted taxonomic labels from all spatial spots (**Methods**) on five taxonomic levels (x axis) when using Kraken2 (orange) or Kraken2 together with the deep learning (DL) model (blue) (y axis) (n = 3). **(c)** Highly specific mapping of bacterial reads. Overall bacterial alignment rates to the respective reference genomes (y axis, %) for GF (left, n = 3), ASF (middle, n = 3) and SPF (right, n = 3) tissue sections using spatial 16S sequencing. **(d)** High reproducibility of bacterial abundances in SPF mouse colons by SHM-Seq. Percentage (y axis) of the top 10 most abundant bacteria genera in each of three independent samples of SPF mouse colons (x axis). **(e)** SHM-seq compares well to 16S rRNA sequencing. Pseudo-bulk abundances of bacteria genera (dot) from SHM-seq (x axis, SPF mice, n = 3) and bulk 16S rRNA sequencing[89] (y axis, SPF mice, n = 3). Top left: Pearson’s *r.* Shaded areas: 95% confidence interval. **(f)** Enzymatic (SHM-seq) extraction of bacterial content agrees with <u>established</u> mechanic extraction. Pseudo-bulk abundances of each bacteria genera (dot) from SHM-seq when bacterial wall permeabilization was performed enzymatically (x axis, SPF mice, n = 3) or by mechanical extraction (y axis, SPF mice, n = 3). Shaded areas: 95% confidence interval. **(g-i)** SHM-Seq agreement with FISH fluorescent signals in ASF mice tissue targeting ASF502 bacteria. **(g)** Distribution (Box plot, normalized signals per region) and individual measurements (scatter plot, mean signal per region and sample (n = 6)) of scaled normalized ASF502 bacterial counts by FISH (y axis, fluorescence intensity with probe targeting ASF502) and SHM-seq (x axis, read counts) in shared morphological regions of interest (MROIs) (colors, **Method)**. Shaded areas: 95% confidence interval. Boxplots: Center black line, median; color-coded box, interquartile range; error bars, 1.5x interquartile range; black dots; outliers. **(h)** Cross-section of an ASF mouse colon with four regions (red rectangle; 1-4) and their **(i)** zoom-in images. Colors: tissue (blue), fibers (gray) and ASF502 bacteria (red).

The deep learning model enhanced performance compared to using Kraken2 alone (**Fig2b, Supplementary Fig1b-d**). First, the deep learning model (when used alone) assigned sequences to genera with 97% accuracy (**Supplementary Fig1b**), on a test dataset of 20% of the data. Next, we used the simulated data to assess taxonomic assignment metrics by comparing the predicted to the true taxonomic labels for data unseen by the model. The deep learning model following Kraken2 outperformed Kraken2 alone significantly on genus level assignment, by several measures, including (1) higher similarity between relative bacterial abundances based on the model’s assignment *vs.* the ground truth (average Pearson’s correlation coefficient for Kraken2 + deep learning model *vs.* Kraken2 alone: 0.97 *vs.* 0.68, p-value≤0.0001, **Fig2b**); (2) higher similarity in bacterial composition, evaluated at a resolution of 1,000 randomly-assigned spatial spots (average Bray-Curtis dissimilarity 0.06 *vs.* 0.15, **Supplementary Fig1c**), and (3) improved total accuracy (92% *vs.* 84%, p-value≤0.0001), F1 score (89% *vs.* 85%, p-value≤0.0001), and lower false positive rate (8% *vs.* 16%, p-value≤0.0001, **Supplementary Fig1d**), when evaluated as bulk-like samples. Thus, the deep learning model can improve the taxonomic assignments.

### Sensitive and specific bacterial rRNA and host mRNA capture

We evaluate SHM-seq by (**1**) specificity and sensitivity of bacterial capture rates compared to bulk 16S rRNA sequencing and by fluorescent *in situ* hybridization (FISH); and (**2**) host RNA-seq quality metrics obtained compared to Spatial Transcriptomics alone.

To assess specificity and sensitivity, we compared profiles from the defined community in ASF[29] mice, as a positive control, and from germ-free (GF) mice, as a negative control (**Fig1**, **Methods**). On average, 22% of all reads in ASF mice samples (n=3) aligned to the bacterial reference, whereas 0% reads from GF mice were assigned as bacterial (n=3) (**Fig2c**), highlighting the specificity of our capture. Bacterial reads mapped to the expected locations in the respective ASF reference genomes (mean reads in expected genomic bin: 97.0±1.5% (mean±SEM, n = 18, **Supplementary Fig2**), with 85.7±4.5% of reads (mean±SEM, n = 18) mapping on average to the expected capture region (**Supplementary Fig3**). Highlighting the sensitivity of SHM-seq, all of the expected bacterial species were captured in the ASF samples, with ASF519 and ASF502 as the dominating bacteria (**Supplementary Fig4a**), in line with previous bulk RT-qPCR results[30] (Pearson’s r = 0.85, **Supplementary Fig4b)**, and with higher reproducibility across replicates (**Supplementary Fig4a**). Even ASF360, which was previously reported to be difficult to detect at low abundances using RT-qPCR[31], was detected with SHM-seq.

**Fig3.**
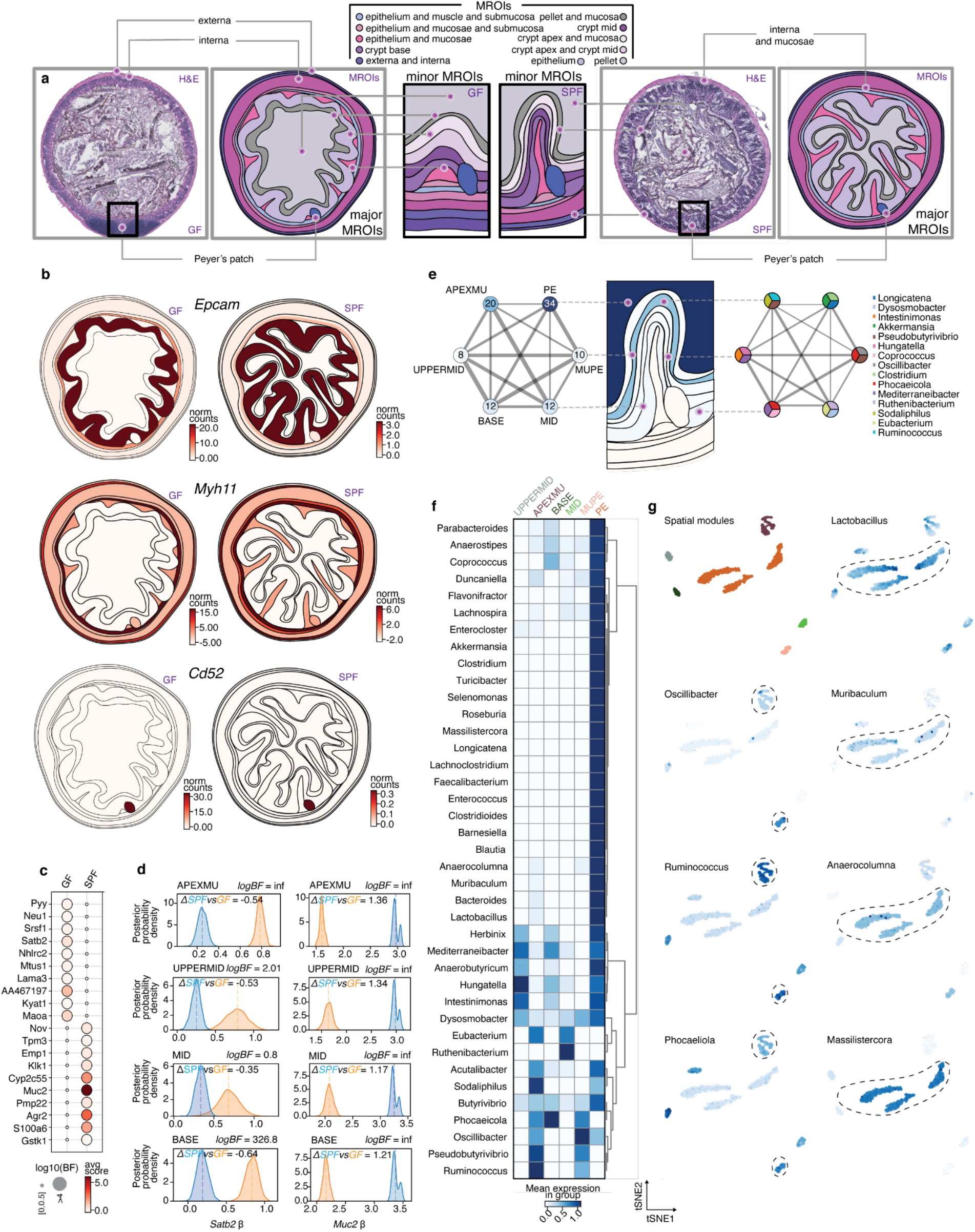
Spatial detection of bacteria and host gene expression with SHM-seq. **(a)** Morphological regions of interest (MROI) in the mouse colon. H&E stained tissue sections from GF (left) and SFP (right) mice (left panels) are annotated and visualized with vector representations (right panels), showing bacteria and host gene expression in major and minor MROI tags associated with each anatomical tissue layer (right panels). **(b)** Spatial host gene expression in three major MROIs. Expression (color bar, normalized gene expression) of selected spatially variable genes in GF (left) and SPF (right) tissue sections in major MROIs. **(c)** Differential gene expression between mouse conditions. Significance (dot size, log_10_(Bayes Factor) (BF), **Methods**) of differential expression and expression level (normalized gene expression) of the top 10 genes (rows) differentially expressed between GF and SPF mouse tissue sections (columns) (**Methods**). **(d)** Gene expression differences between morphological regions. Posterior distributions of the region-specific coefficient parameters (β) of *Satb2* (left) and *Muc2* (right) in four MROIs describing colonic crypts in SPF (blue) and GF (orange) mice. Dashed lines: mean of each distribution. **(e)** Detected bacteria across MROIs in SPF mouse tissues. Number of (left) and top three most abundant (right) bacteria genera (color legend) detected in minor MROIs (middle). Line thickness: Euclidean distances between MROIs. **(f)** Regional abundance of taxa. Scaled normalized bacterial counts (normalized counts scaled within each genus, color bar) in MROIs (columns) for each detected bacteria (rows). **(g)** Association between taxa and spatial regions. t-distributed Stochastic Neighborhood Embedding (t-SNE) of scaled bacterial count profile of each spatial spot (dots) in six minor MROIs in SPF mice, colored by MROI (upper left) or by scaled normalized bacterial counts (color bar as in f) of different genera. MROIs abbreviations (d-f): pellet (PE), mucosa and pellet (MUPE), crypt apex and mucosa (APEXMU), crypt apex and crypt mid (UPPERMID), crypt mid (MID) and crypt base (BASE).

As a more complex case study, we further tested SHM-seq performance in bacterial capture with SPF mice (**Methods**, n = 3). On average, 28% of all reads aligned to the bacterial reference (**Fig2c**), and were assigned to 39 genera in our metagenomic reference (22 of which were present at >1% abundance), with *Duncaniella, Turicibacter* and *Muribaculum* the most abundant (**Fig2d**). The genera detected and their relative abundances correlated well with 16S rRNA sequencing (Pearson’s r = 0.69, p-value≤0.0001, **Fig2e**). Notably, our enzymatic cell permeabilization protocol was as efficient for preparing (bulk) bacterial samples as was traditional mechanical extraction of nucleic acids (Pearson’s r = 0.95, p-value≤0.0001, **Fig2f**).

To further validate the specificity of spatial capture of bacterial genomes in different regions of interest, we compared the bacterial abundance profiles obtained with SHM-seq in ASF mice with those obtained with FISH (**Methods**) with five fluorescent bacterial detection probes: a positive control to detect all bacterial species, probes targeting three distinct ASF species and a negative control. We detected and quantified the fluorescence signal over three major tissue regions (**Methods**, **Supplementary Fig5a-d**). All three ASF specific bacterial abundances correlated significantly between SHM-seq and FISH, including for overall bacterial signals in the positive control probe (average Spearman correlation; ASF502: 0.72, ASF360: 0.72, ASF519: 0.55, positive control: 0.75, p-value≤0.0001, **Fig2g-i, Supplementary Fig5e-g**).

Finally, we confirmed that host RNA-seq quality metrics were comparable between SHM-seq and Spatial Transcriptomics. There were no significant differences in RNA-seq read mapping rate or unique molecular identifier (UMI) counts between Spatial Transcriptomics and SHM-seq in either SPF or ASF mice (n = 3, **Supplementary Fig6a-d**): 66% and 63% of the spatially captured reads were uniquely mapped and pseudo-bulk UMI counts correlated highly (Pearson’s r = 0.95 and 0.92, respectively). Furthermore, there was high agreement in host expression profiles when we used regular ST arrays (only poly(d)T capture) with the permeabilization method developed solely for disrupting host cells *vs*. the one used for disrupting both host and bacterial cells (Pearson’s r = 0.94, **Supplementary Fig6e,f**). Thus, the surface treatment, permeabilization method and library preparation used in SHM-seq compare in specificity and sensitivity to commonly-used methods for accessing bacterial sample composition and for spatial host expression profiling.

### Defining spatial patterns of bacterial organization and host expression

To recover the spatial organization of microbes and host from our data, we defined the expression of host genes and abundance of bacterial genera in each spot, and then mapped those to 16 defined morphological regions of interest (MROIs) (**Fig3a**), to identify characteristic patterns. Briefly, we manually assigned 16 MROIs in each of the profiled tissue sections, leveraging histology guided information, and then used those as rasterized vector representations of tissues for each mouse condition (**Methods**). Overall, we quantified spatial abundances from 100 colonic mouse sections in SPF and GF mice, spanning 10,924 spatially-barcoded spots, each with spatial expression of 17,917 host genes and 39 bacterial genera across the MROIs. On average, we sampled 20 tissue sections, 2,208 spots and ~32,000 nuclear cell segments from each mouse colon (**Supplementary Fig7**). We used a hierarchical probabilistic approach[32,33] (**Methods**) to account for the distance between each sampled spatial spot and MROI, presence of bacteria in the mice (*i.e.* conditions) and individuals (*i.e.* animals) and for differences in sampling (spots) between MROIs, to test for significant spatial expression differences in the sampled sections.

### Spatial co-organization of host expression and microbe composition

We asked how gene expression in each of 16 MROIs was impacted by overall bacterial presence by comparing SPF *vs*. GF mice (with no bacteria). Although both SPF and GF mice showed similar regional expression of some marker genes (*e.g. Epcam* in the epithelium, *Myh11* in the muscularis regions and *Cd52* in Peyer’s patches; **Fig3b**), other genes were significantly differentially expressed between them in particular regions (**Fig3c**). For example, *Satb2* and *Muc2* were, respectively, down- and up-regulated in the crypt apex of SPF *vs*. GF mice, the tissue layer most proximal to the mucosa and lumen (**Fig3d**). *Satb2* helps maintain intestinal homeostasis, and its expression prevents excessive crypt damage and inflammation[34]. Similarly, *Muc2* is key for maintenance of a healthy mucosal layer and its depletion results in direct contact between epithelial cells and bacteria in the colon, leading to inflammation and cancer[35]. In other examples, *Hnf4a,* a gene associated with epithelium renewal[36], is more highly expressed in the base of the crypt in GF *vs*. SPF mice (**Supplementary Fig8**), and *Gpx2,* whose deficiency is related to propagating IBD symptoms[37], is induced in the region bordering epithelium and muscularis mucosae tissue in SPF *vs*. GF mice.

The host spatial expression patterns in SPF mice were mirrored by distinct bacterial genera present at different abundances and compositions in six different MROIs in the SPF mice. The detected bacteria were found in either the colonic interfold regions (crypt base, crypt mid and crypt apex/mid), the mucosal layers (crypt apex/mucosa and mucosa/pellet) or the lumen (*i.e.,* pellet, where they were most abundant, as expected). Interfold regions had the lowest diversity, and the pellet had the highest diversity (**Fig3e**). Morphological regions in close proximity to each other shared some highly abundant genera: *Pseudobutyrivibrio* was shared in the two mucosal regions and *Mediterraneibacter*, an obligate anaerobe and formerly part of the *Ruminococcus* genus[38], was shared between the interfold regions. Mucosal regions had an abundance of *Oscillibacter, Ruminococcus* and *Phocaeliola,* with the latter two genera previously associated with the mucosa[13,39–41], while the pellet had an abundance of commensal bacteria[14,42], such as *Lactobacillus, Muribaculum* and *Anaerocolumna,* but also *Massilistercora,* part of the *Eubacteriales* family and previously reported only in the human gut[43] (**Fig3f,g**).

The mucosal barrier, otherwise preventing unwanted lumen content from coming in direct contact with the host cells in the crypt apex, signals the immune system in a process mediated by epithelial cells[44]. We hypothesized that detected bacteria genera, some observed exclusively with tight junction mucosal barriers (*e.g. Pseudobutyrivibrio, Ruminococcus* and *Oscillibacter,* **Fig3f,g and 4a**) and others observed diffusing into the tissue-specific interfold regions (*e.g. Intestimonas, Coprococcus* and *Flavonifractor,* **Fig3f and 4a**) could influence and be influenced by host expression in close proximity. To systematically investigate significant regional and cell type composition differences and associate them to the presence of bacteria from different genera, we identify 28 gene co-expression modules based on similarity in spatial gene expression (**Supplementary Fig9a, Methods**). We then partitioned each module into sub-modules by gene co-variation across snRNA-seq profiles (**Fig4b, Supplementary Fig9b, Methods**), recovering 203 submodules (**Supplementary Table1, Methods**). We labeled each submodule by its expression in one or multiple of the 30 cell types identified by snRNA-seq and tested it for enriched KEGG pathways (**Fig4c**, **Methods**).

**Fig4.**
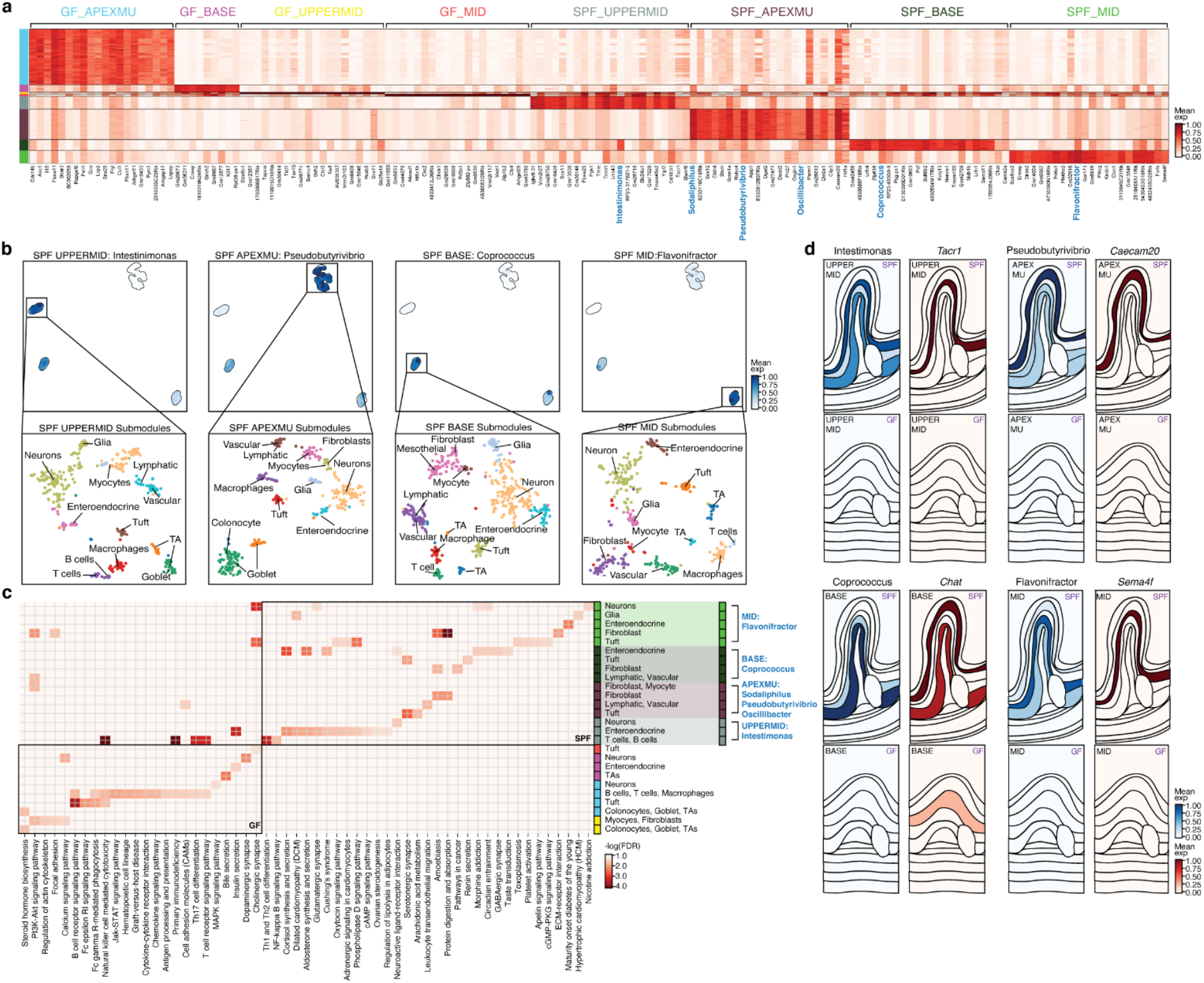
Bacterial presence influences host expression in four major tissue regions. **(a)** Regional association of bacterial taxa and host gene expression. Scaled gene expression (color scale) of top differentially expressed genes (columns, black text) and scaled abundance of top differentially abundant taxa (columns, blue text) in each spatial region (rows) across four major tissue MROIs (color code, left and labels on top). **(b)** Regional association of taxa and cell type composition. Top: t-SNE of scaled bacterial count vectors in each spatial spot (dots) colored by abundance of taxa (color scale) that are differentially abundant in each of four MROIs (labels on top). Bottom: t-SNE of host snRNA-seq cell profiles (dots) mapped in each individual MROI colored by cell type label. **(c)** Expression submodules in different regions reflect distinct biological processes associated with bacterial presence. Significance (color scale, -log_10_(FDR), one-tailed Fischer’s exact test) of enrichment of KEGG pathway genes (columns) in each submodule (rows; black text) associated with each spatial region and mouse condition (color code; right). Blue text: bacterial taxa associated with the same spatial region and condition. Color coding of spatial modules as in **(a)**. **(d)** Differential regional gene expression associated with bacterial taxa in SPF mice. Region maps colored by normalized spatial count of bacterial genera (blue color scale, left panels in panel pair) and normalized spatial expression of cell type marker genes (red color scale, right panels in panel pair) in each MROI.

In the presence of microbiota, specifically *Pseudobutyrivibrio, Sodaliphilus,* and *Oscillibacter,* colonocytes in the apex of the crypts expressed *Caecam20,* a known receptor for Gram-negative bacteria[45] and a known colitis suppressor[46], while goblet cells expressed high levels of *Hif1a,* a marker indicative of a functioning mucosal barrier that is otherwise downregulated in inflammatory bowel disease[47] (**Fig4d, Supplementary Table2**). Neurons in the neighboring region (*i.e.* upper mid region of the crypts), in the presence of *Intestimonas,* expressed *Tacr1* and other neuroactive ligands and receptors implicated in regulating gut motility[48], while macrophages in the same regions and in the presence of the same bacterial genera, expressed *Fcrls* and *Slamf6,* genes that have been shown to modulate neuro-immune signaling upon receptor-microbe binding[49,50] (**Fig4d, Supplementary Table2**). Specialized spatial niches in lower regions of the crypts also contained networks of neurons and myocytes involved in muscle contractility (*Camk2a* in the presence of *Coprococcus),* axon guidance (*Sema4f* in the presence of *Flavonifractor)* and cholinergic signaling (*Chat* in the presence of *Coprococcus)* and (**Fig4d, Supplementary Table2**).

## DISCUSSION

Here, we presented SHM-seq, a method that relies on solid surface capture of polyadenylated host transcripts and variable 16S bacterial regions onto spatially barcoded microarrays for joint spatial profiling of bacterial composition, host gene expression and tissue histology. We provide a deep learning-based approach to enhance taxonomy assignment for metagenomic taxa classification with improved detection rates and assignment accuracy, and a roadmap for interrogating coordinated spatial expression programs.

We benchmark the sensitivity and specificity of SHM-seq compared to traditional 16S sequencing, published RT-qPCR data, as well as FISH and ST in three mouse conditions: SPF, GF and ASF. SHM-seq showed reproducibility and robustness using a tissue dataset of 124 sections and detected all the bacteria genera otherwise present after 16S sequencing in SPF mice as well as all of the 8 species referenced in ASF mice. Previous studies show there is variation in bacterial abundance between mice[30,31,51]. In our study, we also see differences in abundance obtained with SHM-seq *vs.* external ASF data, although the overall correlation between the datasets was high (Pearson’s R: 0.85) and SHM-seq was highly reproducible across mice.

Using these data and methods, we show that in the presence of microbiota, subpopulations of goblet and colonocyte cells formed cell-adhesive layers filled with *Muc2* and *Ceacam1* for host-microbial communication. Additionally, we observed distinct submodules of genes important for a local microenvironment in SPF mice that can regulate intestinal physiological functions and colonic motility, an effect which is known to be disrupted in GF mice[52]. Taken together, these results indicate that our spatial analysis identified expression programs throughout the tissue crosssection characteristic of regional populations that display distinct, mouse condition-relevant dynamics and may depend on the presence of commensal bacteria and/or impact host-bacteria interactions.

SHM-seq enables robust spatial host-microbiome profiling from a large number of tissues, but is currently limited by the resolution of solid-phase capture arrays. Future studies can tackle the resolution limitation using higher density formats[22], and with developing enhanced computational mapping approaches for deconvolving cell-cell interspecies communication networks. Our method paves the way for future work and detailed investigation in larger studies, designed to compare animal models, for example during colitis-induced changes[53] or infection[54], and human patients sampled longitudinally or cross sectionally, where both microbiome and host cells vary, as does host genetics. Such analyses can strengthen the association between host and microbiome features, and lead to better understanding of mechanisms sustaining homeostasis in health, or onset and persistence of chronic inflammation. Our method should thus help in better understanding of environmental and microbiome-driven spatial neighborhood heterogeneity in barrier and mucosal tissues.

## METHODS

### Mice

Adult C57BL/6 specific pathogen-free mice (SPF) were purchased from The Jackson Laboratory (Bar Harbor, ME) and maintained in accordance with ethical guidelines monitored by the Institutional Animal Care and Use Committees (IACUC) established by the Division of Comparative Medicine at the Broad Institute of MIT and Harvard, and consistent with the Guide for Care and Use of Laboratory Animals, National Research Council, 1996 (institutional animal welfare assurance no. A4711-01), with protocol 0122-10-16. Adult C57BL/6 germ-free mice (denoted GF) were obtained from Taconic Biosciences (USA) and maintained in a gnotobiotic environment. Some of these mice were randomly selected and inoculated with ASF[29] over several generations and used when >6 weeks of age. After colonization, ASF mice (denoted ASF) were housed in sterile conditions, and tested with PCR to ensure sterility was maintained[55]. In order to capture material from multiple sections per colonic tube as well as maximize the use of a single spatial array (1,007 spatial spots spread over ~42μm^2^), we placed 2-3 tissue cross-sections onto one spatial capture area. We sampled ~20 sections from each mouse by sectioning in the aforementioned fashion across one spatial capture slide containing 6 active capture areas.

### Tissue collection

Colonic tubes were dissected within minutes of sacrificing mice and tissues dried from excess fluids and embedded in Optimal Cutting Temperature (OCT, Fisher Healthcare, USA) in large molds (VWR, USA) pre-filled with OCT. The molds were then laid onto a metal plate prechilled and set on top of dry ice for 2min or until complete freezing. Samples were transferred to −80°C until sectioning.

### Generation of slides with customized surfaces

Customized surface primers were immobilized to an amine-activated surface area (~40mm^2^ each) using covalent bioconjugation[23,25], as recommended by the manufacturer (Surmodics, USA). Three distinct surfaces were generated for validations: 16S, poly(d)T and a mixed poly(d)T/16S surface. The oligonucleotides immobilization in each case were: 5’-[AmC6]UUUUGACTCGTAATACGACTCACTATAGGGACACGACGCTCTTCCGATCTNN NNNNNNATCTCGACGACTACHVGGGTATCTAATCC-3’ 5’-[AmC6]UUUUUGACTCGTAATACGACTCACTATAGGGACACGACGCTCTTCCGATCTN NNNNNNNTTTTTTTTTTTTTTTTTTTVN-3’ (both IDT, USA).

All slide incubations took place on a thermal incubator (Eppendorf Thermomixer Option C, Germany) with slides mounted into a hybridization chamber (ArrayIT, USA). All *in situ* reactions performed on spatial arrays were carried out in biosafety cabinet class II.

### Generation of spatial arrays with customized surfaces

All spatial arrays were produced as previously described[23,25]. Briefly, six spatial microarrays were created on amine-activated Codelink slides (Surmodics, USA). Printing was performed by ArrayJet LTD (Scotland, UK) spotting spatially barcoded DNA oligonucleotides as 100μm spots. A total of 1,007 different oligonucleotide addressable spots was printed per capture area. To hybridize the 16S probe onto the surface probes of a spatial array, 75μl of a mix of the 16S probe (5’-GGATTAGATACCCBDGTAGTCGAGATNBAAAAAAAAAAAAAAAAAAAA-3’, IDT, USA) at 0.6μM concentration with 4X Saline Sodium Citrate (SSC) (SSC, Sigma-Aldrich, USA) and 0.1% SDS (Sigma-Aldrich, USA) was added to each well and incubated for 2h at room temperature (RT). The probe mix was then removed and the wells were washed with 100μl of 0.1X SSC (SSC, Sigma-Aldrich, USA). To covalently attach the now hybridized 16S probes onto the surface probes, 75μl of 0.5mM dNTPs (Thermo Fisher Scientific, USA), 5U Klenow Fragment (3’→5’ exo-), 1X NEB Buffer 2 and 0.20μg/μl BSA (all from NEB, USA) was added to the wells and incubated at 25°C for 1.5h. The solution was then removed, followed by a wash with 100μl 0.1X SSC. To strip off the 16S probes and make surface probes single stranded again, wells were 3X washed with 75μl 35% formamide for 30sec at RT. Wells were then washed twice with 100μl 0.1X SSC, before slides were detached from their hybridization chambers and washed for 10min at 50°C in 2X SSC/0.1% SDS (Sigma-Aldrich, USA), followed by 1 minute wash with 0.2X SSC and finally 0.1X SSC, both at 37°C. Slides were dried and immediately used in cryosectioning.

### Cryosectioning

The entire cryo chamber including all surfaces and tools used during cryosectioning were wiped with 70% EtOH before the start of sectioning to avoid bacterial contamination. Both spatial arrays and OCT-embedded gut tissue blocks were allowed to reach the temperature of the cryo chamber, before 10μm thick cross-sections of gut tissue were placed on customized spatial arrays. Tissue fixation followed immediately as described below.

### Tissue fixation, H&E staining and imaging

The spatial array was warmed at 37°C for 1min. Then, the entire area of the glass slide was covered in a methacarn solution (60% absolute methanol, 30% chloroform stabilized with ethanol and 10% glacial acetic acid (all from Sigma-Aldrich, USA)) for 10min at RT in a closed space to avoid evaporation. Methacarn was then removed and the slide was allowed to dry before ~300μl of isopropanol (Sigma-Aldrich, USA) were added to the slide and incubated for 1 minute at RT. When the slide was completely dry again, it was stained using hematoxylin and eosin (H&E) staining in an EasyDip Slide Jar Staining system (Weber Scientific, USA). The system included containers separately filled with ~80ml Dako Mayer’s hematoxylin, Dako Blueing buffer (both from Agilent, USA), 5% Eosin Y in 0.45M Tris acetate (both from Sigma-Aldrich, USA) buffer at pH 6 and nuclease-free water (ThermoFisher Scientific, USA). The slide was put in a slide holder and completely dipped in hematoxylin for 8min, followed by five dips in nuclease-free water and then ten dips in a beaker filled with ~800ml nuclease-free water. The slide holder was then dipped in Dako blueing buffer for 1min, followed by another five dips in nuclease-free water. Finally, the slide holder was put in the eosin solution for 1min, and washed by five dips in nuclease-free water. The slide was removed from the holder and air dried before mounted with 85% glycerol and covered with a cover slip (VWR, USA) prior to imaging. Imaging of stained H&E tissue sections on glass arrays was performed on a Metafer Vslide scanning system (MetaSystems, Germany) installed on an Axio Imager Z2 microscope (Carl Zeiss, Germany) with a LED transmitted light source and a CCD camera. Using an A-P 10×/0.25 Ph1 objective lens (Carl Zeiss, Germany) and a configuration program[24], focusing and scanning of each tissue section on the glass array was done automatically. Image stitching was done using Vslide (v.1.0.0) with 60μm overlap and linear blending between fields of views. Images were extracted using jpg compression.

### *In situ* reactions: permeabilization and reverse transcription

Before start, the hybridization chamber was cleaned with RNaseZap (Fisher Scientific, USA) and 70% EtOH followed by at least 30min in a UV light chamber. After section imaging, the slide was again attached to the hybridization chamber to proceed with the following permeabilization reactions (referred to as: “bacterial treatment” below). First, 100μl of a lysozyme solution with 200ug/μl lysozyme (Sigma-Aldrich, USA) was added to each well and incubated for 1h at 25°C, followed by wash with 100μl 0.1X SSC. Second, 75μl of 0.01X Triton X-100 (Sigma-Aldrich, USA) was added and incubated for 15min at 25°C, followed by a 100μl wash of 0.1X SSC. Lastly, 75μl per well of 0.1% pepsin (pH 1, Sigma-Aldrich, USA) was incubated for 7min at 37°C, followed by a final wash of 100μl 0.1X SSC. *In situ* cDNA synthesis was performed as previously described[24]. Briefly, 75μl of 50ng/μl actinomycin D (Sigma-Aldrich, USA), 0.5mM dNTPs (Thermo Fisher Scientific, USA), 0.20μg/μl BSA, 1 U/μl USER enzyme (both from NEB, USA), 6% v/v lymphoprep (STEMCELL Technologies, Canada), 1M betaine (#B0300-1VL, Sigma-Aldrich, USA), 1X First strand buffer, 5mM DTT, 2U/μl RNaseOUT, 20U/μl Superscript III (all from Thermo Fisher Scientific, USA) were added to each well. The reaction was sealed with Microseal ‘B’ PCR Plate Seals (Biorad, CA) and incubated for at least 6h. After incubation, 70μl of the released cDNA material from each hybridization chamber well were collected and stored in a 96-well PCR plate (Eppendorf, Germany).

### Library preparation

Library preparation was performed using the SM-Omics automated library preparation protocol, as previously described[24]. Briefly, released cDNA material was first made doublestranded using the nicked RNA template strands as primers to copying the cDNA strand with DNA polymerase I. To avoid overdigestion, the reaction was terminated with EDTA and ends were blunted using T4 DNA polymerase, before linear amplification by *in vitro* transcription. Amplified material was again transcribed into cDNA resulting in material ready for PCR indexing as described in the next section.

### Quantification, indexing and sequencing

qPCR quantification and indexing were performed as previously described[56] using TruSeq LT Illumina indexing and a KAPA HotStart Hifi Ready-Mix (Roche, Switzerland). Indexed cDNA libraries were cleaned using a 0.7:1 ratio with Ampure XP beads (Beckman Coulter, USA) to PCR product, according to the manufacturer’s protocol and eluted in 12μl elution buffer (Qiagen, Germany). Each sample’s concentration was measured using the DNA HS Qubit assay (ThermoFisher Scientific, USA) and average fragment length was determined using either Bioanalyzer HS or DNA1000 Tapestation (both from Agilent Technologies, USA). Each sample was then diluted to the desired concentration for sequencing (1.08pM on a NextSeq and 10pM on a Miseq, both with ~10% PhiX). Pooled libraries were sequenced with 25nt in the forward read and 55nt and 150nt in the reverse read on NextSeq and MiSeq (Illumina), respectively.

### Mechanical extraction of bacterial RNA

Approximately 1mm thick tissue section with pellet was sectioned from SPF colons in OCT and put in dry ice-cold Lysis matrix D tube (MP Biomedicals, USA). 400μl of RLT buffer (Qiagen, Germany) with 1% 2-Mercaptoethanol (Sigma-Aldrich, USA) were added to the tube and the solution was homogenized in a FastPrep-24 instrument (MP Biomedicals) at speed 6 for 40 seconds. Tubes were then centrifuged for 5min at 12,000 rpm. Supernatant was transferred to a new tube and RNA extraction was done using the RNeasy mini kit (Qiagen), according to the manufacturer’s instructions. Extracted RNA was fragmented using the NEBNext Magnesium RNA Fragmentation module kit (NEB, USA), heating for 2min. Fragmented RNA was cleaned with the MinElute Cleanup kit (Qiagen), according to the manufacturer’s instructions. Quality of the fragmented RNA was evaluated by the Bioanalyzer pico kit (Agilent Technologies, USA). ~20ng/μl of mechanical extracted RNA were added on a 16S surface probe coated QC array in an *in situ* cDNA reaction, as described in the *“in situ reactions: permeabilization and reverse transcription”* section. After at least 6h incubation at 42°C, 70μl of the released material from each well were collected and stored in a new 96-well PCR plate (Eppendorf, Germany). Library preparation, quantification, indexing and sequencing on the MiSeq were performed as described in the *“Library preparation* and *Quantification, indexing and sequencing”* section.

### Extraction and metagenomic sequencing of fecal DNA

Pellet was collected from the colon of SPF mice by perforating the colon wall and scraping the pellet and mucus into a 1.5 mL collection tube (Eppendorf, Germany). Collected pellet was stored at −80°C until further processed. DNA was extracted from the pellet using a Lysing matrix Y tube (MP Biomedicals, USA), according to the manufacturer’s instructions. Extracted DNA concentration was determined using the DNA HS Qubit assay. DNA was made into libraries using Nextera XT (v.15031942 v05). Concentration and average fragment length of each sample were evaluated using the DNA HS Qubit assay (ThermoFisher Scientific, USA) and BioAnalyzer HS (Agilent Technologies, USA), respectively. Each sample was diluted to the desired concentration for sequencing (9pM, ~10% PhiX), and pooled samples were sequenced on MiSeq (2×150bp, lllumina). Each sample was sequenced to ~5-10 million reads.

### Raw reads processing and mapping of host reads

Reads were generated with bcl2fastq2 (v.2.20.0) and trimmed to remove adaptor sequences and the 16S surface probe sequence using BBDuk[57] (v.38.33). ST Pipeline (v.1.7.6)[27] was used to generate gene-by-barcode matrices. The reverse quality-filtered reads were mapped with STAR (v2.6.0)[58] to the mouse genome reference (GRCm38 primary assembly), and mitochondrial sequences were removed. Mapped reads were annotated using HTseq-count (v0.11.4)[59] and the mm11 mouse annotation reference (https://www.gencodegenes.org/mouse/release_M11.html). Annotated reads were demultiplexed with TagGD[27,60] (v0.3.6) with a hamming distance clustering approach (k-mer 6, mismatches 2). This connected transcript information to spatial barcodes. Finally, UMI collapsing per transcript and spatial barcode was performed with a naive clustering approach (mismatches 1) similar to that described in UMI-tools[61].

### Generation of mouse gut bacterial reference

Fastq reads were generated with bcl2fastq2 and reads were quality filtered using KneadData (v.0.7.4) (https://huttenhower.sph.harvard.edu/kneaddata/) (mouse database mouse_C57BL). MEGAHIT[62] (v.1.2.9) was used for assembly of the filtered reads and bowtie2[63] (v.2.3.4.3) for mapping reads to the assembly. MetaBAT2[64] (v.2.15) was used for binning the assembly and the command-line version of NCBI BLAST[65] (v.2.9.0+) was used to assign taxonomy to contigs with blastn and database ‘nt’. MEGAHIT, bowtie2 and MetaBAT2 were all run using default settings. Assignments were filtered (E-value≤10E-6) and sorted (by E-value and percent identity) and each contig was then assigned the top taxonomy assignment. Only contigs belonging to an assigned taxonomy on species level that represented >0.1% were retained. Reference genomes for these species were downloaded from NCBI RefSeq database[66], resulting in a fasta sequence database of the taxa found in SPF mice (n=6) and used as input to build a custom database in Kraken2 (v.2.0.9)[28] according to Kraken2 default instructions, including masking of low-complexity regions. Reference genomes for five species were not found in the RefSeq database and were not included in the fasta sequence database: *Hungatella xylanolytica, Christensenella massiliensis, [Eubacterium] rectale, [Eubacterium] siraeum* and *[Clostridium] innocuum.* The mouse gut bacterial reference was also filtered for genera that have previously been found in mice and/or the intestine[67–69]. A phylogenetic tree of the reference taxa was built using NCBI’s Common Tree and visualized using iTOL (v.6.4.3)[70]. When analyzing mouse gut tissue with defined flora (ASF), genome sequences according to Wannemuehler *et al[71]* were downloaded from NCBI and used as input to build a custom ASF database in Kraken2.

### Generation of simulated data

16S rRNA fasta sequences for the taxa found in SPF mice (determined as described in *“Extraction of fecal DNA and generation of mouse gut bacterial reference”)* were downloaded from NCBI (downloaded 2021-07-24), except two taxa where the 16S rRNA fasta sequence were missing (*Sodaliphilus pleomorphus* and *Anaerocolumna sedimenticola).* Command-line NCBI BLAST[65] (v.2.9.0+) was used to align every possible sequence version of the 16S surface probe to the 16S rRNA fasta sequences to find the best possible alignment for the 16S surface probe per taxa. To mimic spatially captured reads from a real SHM-seq, two million paired reads from a real SHM-seq experiment were used as a template for fastq headers, sequence and quality scores for the forward read and fastq headers and quality scores for the reverse read. The sequences in read 2 were replaced by 150bp long fragments of the 16S rRNA sequences from randomly selected taxa. Fragments were created by selecting a region upstream of the best possible alignment of the 16S surface probe per randomly selected taxa. Each region was then randomly selected a length based on a normal length distribution with parameters characteristic to a spatial array (400 +/− 44bp) and trimmed to 150bp. This resulted in a simulated data set with two million randomly selected 16S rRNA gene sequences, generated from where the 16S surface probe is expected to capture, from the taxa in our mouse gut bacterial reference, but with known exact taxa and both reverse and forward reads.

### Deep learning model: data pre-processing

A total of 500,000 DNA sequences were randomly selected from the simulated dataset (described in *“Generation of simulated data set*”), uniformly sampled, and single-point mutations with 0.1% rate were introduced. This was followed by random shortening, based on a normal distribution of fragment lengths from a true SHM-seq experiment (143 +/− 13bp, truncated at 150bp). Reads from each taxon in the mouse gut bacterial reference were represented at least 100 times per genus. Sequences were one-hot encoded, such that each nucleotide (A, C, T, G and N) was represented by a five-dimensional binary vector, followed by sequence padding up to the maximum length (150bp). Taxa labels were one-hot encoded into one of N genera. The encoded sequences and taxa labels were provided as input for training the model.

### Deep learning model: architecture

A taxonomic classifier of short reads was implemented using Keras[72] with TensorFlow[73] backend (v2.2.0) in Python (v.3.8.10) (**Supplementary Fig1**). The model takes as input one-hot encoded DNA sequences of varying lengths and provides a genus label as output. First, a masking layer was used to ignore padded entries, followed by four layers of a one-dimensional convolutional layer with kernel sizes of 15, 17, 19 and 23 to extract short motifs, followed by a concatenation and a dropout (50% rate) module and two bidirectional long short term memory networks layers, which processed the sequences in both directions. This was followed by another dropout layer (20% rate), a dense layer (reLU activation), a dropout layer (10% rate), another dense layer (reLU activation) and finally a fully-connected layer (softmax activation) to reduce the final output size to the number of distinct genera in the input data. In total, the model consisted of 298,760 trainable parameters. Cross-entropy loss was used to train a multiclass classifier with ADAM as the optimization algorithm[74]. The model architecture was visualized using Netron[75].

### Deep learning model: training details

Model parameters were optimized by using 80% of sequences for training and 20% for testing. Each epoch started with shuffling the training data and computing the gradient update once for each training data point to obtain unbiased gradient estimates[76]. During training categorical accuracy and cross-entropy loss were used to monitor progress. Training was terminated after a maximum of 15 epochs or when the training loss did not decrease in five consecutive epochs. The area under receiver-operating characteristics (ROC) and the F1 score were calculated using Scikit-learn (v.0.24.2)[77] and used to report the final performance on test data.

### Deep learning model: evaluation

One million simulated sequences with corresponding taxa (as in *“Generation of simulated data set”)* were modified with a sequencing error rate of 1%[78] and random shortening as described above. Sequences were classified either by Kraken2 alone or by Kraken2 followed by the deep learning model. Performance was evaluated compared to the ground truth taxa labels by calculating Bray-Curtis distances and Pearson correlation coefficients of the bacterial relative abundances per spot using Scipy (v1.1.0)[79] spatial.distance.braycurtis and Scikit-learn (v.0.24.2)[77] stats.pearsonr, respectively. A higher similarity of the relative abundances between classifications and the ground truth resulted in lower Bray-Curtis dissimilarities and higher Pearson correlations. Accuracy and F1 score were calculated on the whole data set using Scikit-learn (v.0.24.2)[77] metrics.classification_report.

### Raw reads processing and mapping of bacterial reads using taxonomy assignment pipeline

Fastq reads were generated with bcl2fastq2 and trimmed to remove adaptor sequences using BBDuk[57]. Trimmed reads were quality filtered using the same quality filtering step as in ST pipeline (v.1.7.6)[27] but only reads longer than 100nt were kept. TagGD[60] was used to connect the spatial barcode to each forward read (k-mer 6, mismatches 2, hamming distance clustering algorithm) and BWA mem (v0.7.17)[80] with reference mouse genome (GRCm39) was used to remove host mapping sequences. Remaining reverse reads were mapped to the mouse gut bacterial reference using Kraken2 (v.2.0.9)[28] (confidence 0.01). Taxonomy assignments made by Kraken2 were improved using the deep learning model. UMIs with identical spatial barcodes and taxonomical assignments were collapsed using UMI-tools (v.1.0.0)[61] (UMIClusterer, threshold 1), resulting in a bacteria-by-barcode matrix.

### Fluorescence *in situ* hybridization

FISH was performed on the same fresh frozen gut tissue samples from ASF mice. All sections were 10μm thick cross sections and consecutively collected; first sections were placed on the spatial array, followed by placing consecutive sections on a Codelink amine-activated slide (Surmodics, USA), the following two sections were then again placed on the spatial array. Sections on the spatial array were used for SHM-seq and sections on the amine-activated Codelink slide (Surmodics, USA) were prepared for FISH as further described. Slides were warmed at 37°C for 2.5min on a thermal incubator, before tissue sections were fixed using freshly prepared methacarn, as described in *“Tissue fixation, H&E staining and imaging”.* Slides were then placed in a hybridization chamber, and 75μl of preheated FISH solution (0.9M NaCl, 20mM Tris pH 7 (both Thermo Fisher Scientific, USA), 0.1% SDS (Sigma-Aldrich, USA)), a FISH oligonucleotide detection probe (0.06ug/ul)) were added to each well and incubated for two hours at 25°C. Oligonucleotide detection FISH probes (IDT, USA) were used depending on the target of interest: probe EUB338 (5’-/Cy5/GCTGCCTCCCGTAGGAGT-3’) for all bacteria, probe non338 (5’-/Cy5/ACTCCTACGGGAGGCAGC-3’) as a negative control, probe Lab158 (5’-/Cy5/GGTATTAGCAYCTGTTTCCA-3’)[81–83] to target ASF360, probe Lac435 (5’-/Cy5/TCTTCCCTGCTGATAGA-3’)[84,85] to target ASF502 and probe Bac303 (5’-/Cy5/CCAATGTGGGGGACCTT-3’)[8,83] to target ASF519. After the two hours incubation, FISH solution was removed and wells were washed with 100μl 1X PBS, before the hybridization chamber was removed and slides were dipped 12 times in 50ml 1x PBS before air dried. Slides were mounted with 85% glycerol (Sigma-Aldrich, USA) and a cover slip (VWR, USA). Epifluorescent images were acquired on an Axio Imager Z2 microscope using a PhotoFLuor LM-75 lightsource (89North, USA) in combination with a Plan-APOCHROMAT 63x/1.4 Oil DIC objective (Carl Zeiss, Germany). Images were processed using Vslide (v1.0.0, MetaSystems GmbH).

### Spatial analysis of bacterial fluorescence

Bacterial presence in scanned fluorescence images was detected using Ilastik (v1.3.3)[86]. After training and testing each bacterial fluorescence print separately in Ilastik, the resulting bacterial detection mask was aligned with the fluorescence image to detect mean fluorescence intensity per spatial coordinate and stored as a matrix. This matrix was then run in Splotch (as described in *“Spatial modeling of host-microbiome data”*). Resulting normalized fluorescence intensity was compared to the normalized bacterial presence by randomly selecting at most three spatial coordinates from each annotated region per sample (only annotated regions that were shared between the normalized fluorescence intensity and the normalized bacterial presence were considered) and scaling them within each sample, before matching them to a spatial coordinate in the same region and comparing them to each other (normalized fluorescence intensity *vs.* normalized bacterial presence per spatial coordinate). To limit the region annotated as pellet, spatial coordinates annotated as pellet were selected if they were spatially adjacent to coordinates annotated as mouse tissue. This procedure was repeated a thousand times to generate an average spatial correlation measurement between normalized bacterial FISH intensity and normalized sequenced bacterial presence, expressed as Spearman correlation.

### 16S surface probe sensitivity

To evaluate 16S surface probe sensitivity, reference DNA sequence and gene annotation files were downloaded from Ensembl Bacteria[87] for the ASF bacteria available in the database (v104.1) (ASF356, ASF360, ASF457, ASF492, ASF500 and ASF519 (taxonomy id 1235789)). Reads captured from ASF tissue sections on a ST QC array with only 16S surface probes on the array surface were separately mapped against each ASF bacteria genome using BWA mem (v0.7.17)[80]. Gene body coverage over the 16S rRNA genes in respective reference genomes was generated using RSeQC (v4.0.0)[88]. Genome binning was done by summarizing the aligned reads in separate bins, each bin representing a hundredth of the respective ASF genome.

### 16S surface probe specificity

Specificity was first evaluated by proportion of bacterial *vs.* mouse read alignment. Tissue sections from SPF, ASF and GF mice were placed on QC arrays with 16S surface probes and finished libraries were prepared using either bacterial treatment or colon treatment. Each finished library was sequenced to approximately 660,000 reads. Reads were taxonomically annotated by using the taxonomy assignment pipeline without the deep learning model. The proportion of reads mapping to the respective bacterial reference (mouse gut bacterial reference for SPF and GF tissue samples and ASF reference for ASF tissue samples) was calculated by using the number of trimmed reads.

Protocol specificity was also evaluated by comparing the bacterial treatment with a mechanical treatment (*“Mechanical extraction of bacterial RNA”).* Spearman’s rank and Pearson correlation coefficients were calculated using Scipy’s (v1.1.0)[79] stats.spearmanr and stats.pearsonr.

Bacterial treatment was compared to a bulk 16S rRNA sequencing dataset[89] where the 16S libraries were made from material originating from feces of C57BL/6J mice (SRA sample references: SRR9212951, SRR9213178, SRR9213335). The correlation was calculated using Scipy’s (v1.1.0) Pearson’s correlation coefficient[79].

### Effect of the bacterial treatment on mouse gene expression

To evaluate the effect of the bacterial treatment on measured host (mouse) gene expression, we normalized[90] gene counts from samples with and without bacterial treatment (reads downsampled to the same saturation levels) and from samples prepared on a spatial array with customized surface or a standard spatial array (reads downsampled to the same saturation levels). Pearson correlation coefficient was calculated using Scipy’s (v1.1.0)[79] stats.pearsonr.

### Image registration and annotation

Image processing and registration of barcoded spots was done using SpoTteR[24]. H&E images (collected in RGB channels) were downscaled to approximately 500×500 pixels. For efficient grid spot detection, tissues were masked from the images using quantile thresholding in the red channel. Centroids of spatial array spots were detected by computing the image Hessian. Centroid coordinates were used as probable grid points and a rectangular grid was then fitted to these probable points using a local optimizer (*nlminb,* R package stats (R v.3.6.3)). With iterations and removing 10% of the probable spots that did not fit the perfect grid structure, a new grid was fitted until the target number of grid points per row (here 35) and column (here 33) were reached. Final grid points were overlapped with the previously masked tissue section to select spatial points only present under the detected tissue section area. These points were used in further analysis.

H&E images were annotated using a graphical cloud-based interface[22] by manually assigning each spatial coordinate (x,y), resulting from the grid fitting process, with one or more morphological region tags. The tags used were epithelium (E), epithelium and muscle and submucosa (ALL), epithelium and mucosae and submucosa (EMMSUB), epithelium and mucosae (EMM), muscle and submocosa (MSUB), crypt base (BASE), externa and interna (MEI), externa (ME), interna (MI), mucosae and interna (MMI), mucosa and pellet (MUPE), crypt mid (MID), crypt apex and mucosa (APEXMU), crypt apex and crypt mid (UPPERMID), Peyer’s patch (PP) and pellet (PE). ALL, E, EMMSUB, EMM, BASE, MEI, ME, MI, MUPE, MID, MMI, APEXMU, UPPERMID, PP and PE were visualized in tissue vector representations.

### Spatial modeling of host-microbiome data

Splotch[32,33] was used for statistical analysis of spatial data. Splotch is a hierarchical probabilistic that captures variation in ST data through modeling of different study design covariates such as individual’s age or mouse condition (*B*), a linear model component capturing spatial variation in array data with a conditional autoregressive (CAR) prior (*ψ*), and gene expression variation captured in each independent spatial measurement (*ϵ*) to account for technical artifacts. Sequencing depth is accounted for by using a size factor *s* where the total number of captured UMI counts per spatial spot is divided by the median UMI counts across all analyzed spots. The posterior distribution of the parameters is interrogated from the model, for example, when the model was conditioned of bacterial presence in the tissues in order to quantitate expression changes across both the mouse conditions and different tissue contexts.

Genes (*i*), tissue sections (*j*), and independent spatial spots (*k*) were indexed as follows: 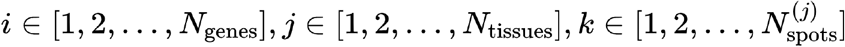. Gene expression in each spot is considered an approximation of observed counts *y_i,j,k_*, where *y_i,j,k_* is expected to equal to *S_j,k_λ_i,j,k_ s_j,k_* is the size factor (total number of UMIs observed at spot *k* and tissue section *j*), and *λ_i,j,k_* is the rate of gene expression (referred to as normalized counts throughout). Splotch then models the observed counts using the zero-inflated Poisson (ZIP) distribution:

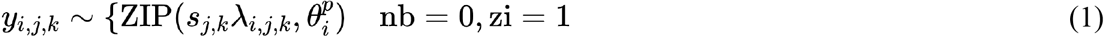

where 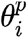 represents the gene-specific probability of a dropout. The zero-inflated models account for an overabundance of zeros by introducing a second zero-generating process gated by a Bernoulli random variable:

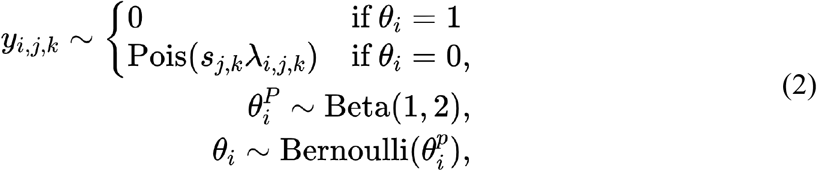

where the Poisson process can be replaced by NB without loss of generality. The gene expression rate parameter *λ_i,j,k_* is described in terms of a generalized linear model (GLM) by three components:

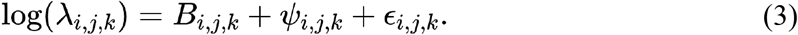

where *B_i,j,k_* is the characteristic expression of gene *i* within the context of spot *k* from which a characteristic expression vector *β_i_* ∈ ℝ^N_MROI_^ is derived describing which MROI spot *k* comes from. At the top level, the dataset is split along an important covariate (*e.g.,* presence of bacteria), and a separate *β_i_,l*_1_ is modeled for each unique group (*l*_1_ ∈ {1,...*L*_1_}). At the next level, each set is further partitioned along another covariate (e.g., animal individual). A two-level hierarchical model for *β_i_* can thus be specified as:

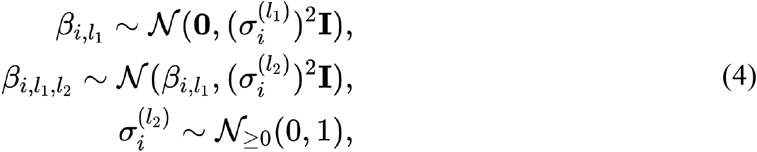

where in practice 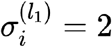 for all *i,l*_1_ and posteriors are inferred over all 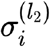. For convenience, because each tissue *j* belongs to one covariate group at each level, the inverse mapping function *ρ*^1^(*j*) is introduced that maps *j* to the appropriate *l*_1_,*l*_2_,*l*_3_ indices for *β_i_*. With this in hand, *B_i,j,k_* is formally defined in the non-compositional model:

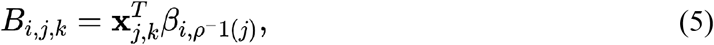

where **x**_*i,k*_ is a one-hot encoding of the spot MROI annotation 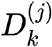 used to index the relevant entry in the characteristic expression vector *β*_*i,ρ*^-1^(*j*)_.

*ψ_i,j,k_* describes the how the local and immediate neighborhood of spot *k* has an effect gene *i* and is modeled using the CAR prior. The observations in each spatial spot are assumed to be dependent on the spot’s immediate spatial neighborhood defined as 4 nearest neighbors. *ψ_i,j_* is defined as a Markov random field over the spots in each array:

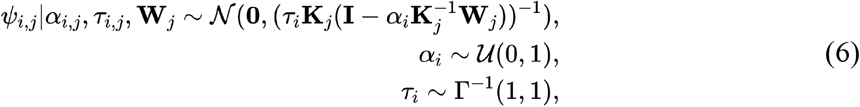

where *α_i_* is a spatial autocorrelation parameter, *τ_i_* is a conditional precision parameter, **K**_*j*_ is a diagonal matrix containing the number of neighbors for each spot in tissue *j*, and **W**_*j*_ is the adjacency matrix (with zero diagonal).

*ϵ_i,j,k_* captures variation at the level of individual spots with the assumption that each spot was independently and identically distributed (i.i.d) in order to infer their standard deviations:

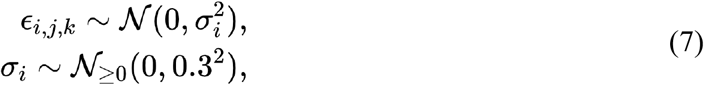

where *σ_i_* is the inferred level of variability for gene *i*.

Data were processed as a two-level model when describing differences in mouse model/condition and morphological region (when comparing SPF *vs*. GF mice), or as a one-level model for ASF mouse analysis. Input data were raw UMI counts (as described above). Sampling from the posterior was done running four independent chains with 200 iterations per chain (100 warmup and 100 sampling). The model was conditioned on 10,924 spots, 16 morphological region tags and 2 mouse conditions (SPF *vs.* GF) (two level model) or 4,821 spots, 5 morphological region tags and one mouse condition (ASF) (one level model).

For differential expression analysis, each pairwise comparison of gene expression was denoted as a random variable Δ_β_, that describes the difference between two conditions as *β*_1_ – *β*_2_. *β*_1_ and *β*_2_ represent any two conditions arising from any two combinations in the model, for example, any two genes, sample covariates (e.g. mouse condition; eg SPF *vs.* GF) or MROIs regions (*e.g.* crypt apex and mucosa *vs.* crypt base). The null hypothesis presumes that the two posterior distributions over characteristic expression coefficients *β*_1_ and *β*_2_ estimated by the model are identical and that **Δ**_*β*_ is tightly centered around zero. To quantify this similarity, 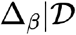 (where 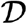 is the training data) is compared to the prior distribution **Δ**_*β*_ using the Savage-Dickey density ratio[91] that estimates the Bayes factor (BF) between the conditions:

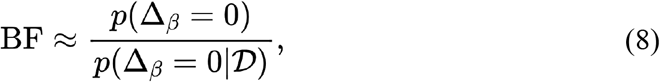

where the probability density functions are evaluated at zero. If expression is different between the two conditions, then the posterior 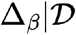 will not be centered around zero, and the estimated BF will be large, hence, the null hypothesis is rejected and the two genes are denoted as differentially expressed between the conditions. Hereinafter, the Savage-Dickey density ratio is referred to as BF. Upregulated genes (**Δ**β > 0) with at least *log*(*BF*) > 0.5 were considered as differentially expressed between any two conditions and used in all downstream analysis. Bacterial genera were called as detected in SPF tissue if the bacterial weighted mean count per morphological region was greater than the maximal weighted mean in corresponding morphological mouse region in GF. The total regional count had to count for more than 2% of the total bacterial count to be called as detected.

### Single-nucleus RNA-Seq data analysis

Mouse colon single-nucleus RNA-Seq (snRNA-seq) data were obtained from Drokhlyansky *et al [92]*, containing 340,461 individual cell profiles across 22,986 expressed genes. Briefly, nucleus profiles with >800 genes expressed in a minimum of 10 cells and <30% mitochondrial or rRNA signatures were retained for analysis. Raw counts data were normalized to transcripts-per-10,000 (TP10K). To regress out genes as differentially expressed, the mean and the coefficient of variation (CV) of expression of each gene were calculated and partitioned into 20 equal-frequency bins. LOESS regression was used to estimate the relationship between log(CV) and log(mean) and genes with the 1,500 highest residuals were equally sampled across these bins. To account for differences in batches, this was performed for each sample separately and a consensus list of 1,500 genes with greatest recovery rates was selected. Next, using Scanpy[93], Harmony[94] was used for further batch correction with 20 neighbors and 40 principal components from PCA. After 10 iterations, convergence was reached and the resulting data clustered with Phenograph[95], with 25 nearest neighbors using the Minkowski metric. Cell type labels provided in Drokhlyansky *et al[92]* were used to manually label clusters after Phenograph clustering.

### Spatial co-expression analysis

All posterior estimates that account for both morphological differences and differences in mouse conditions were used as *λ_i,j,k_* in a sparse matrix format 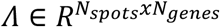, where *N_spots_* = 5,413 and *N_genes_* = 17,956. The snRNA-seq normalized counts and SHM-seq posterior means counts tables were standardized separately across cells and spots within genes, respectively, considering common genes (*N_Common genes_* = 16,525) in both datasets, resulting in matrices 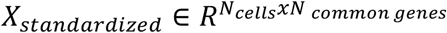 and 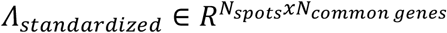. Finally, the similarity of each cell to each spot was calculated as the Pearson correlation coefficient *r* between its standardized and imputed expression vector (columns of *X_standardized_*) and spots’ expression vectors (columns of *Λ_standardized_*), resulting cell-specific similarity vectors (**Supplementary Fig9**). Morphological spots were used from all region categories except for those found in PE and MUPE. To find sets of co-expressed genes *i.e.* with similar spatial patterns, the data *P* were hierarchically clustered with the average linkage method using the *L1* norm (Manhattan distance), with a set distance threshold to detect 28 distinct blocks (hereinafter modules) using scipy.cluster.hierarchy.fcluster.

### Using snRNA-seq profiles to partition spatial clusters to submodules

Gene expression submatrices were created of the expression of genes belonging to each co-expression module. To identify which specific cell types underlie expression in each spatial module mean expression values were calculated for each gene across the single cell profiles in each of 30 snRNA-seq clusters (as described in “*Single-nucleus RNA-Seq data analysis”*) and scaled by dividing each gene’s mean expression per cluster by its maximum mean expression across cell-type clusters. Genes with an average scaled expression lower than 1 were removed (scaled expressions set to zero). Then, to estimate cell type compositions in each module, these filtered and averaged snRNA-seq signatures were hierarchical sub-clustered within each module using cosine distance and average linkage. Genes were grouped in submodules using 0.4x the maximum of the linkage matrix as cutoff. Two-sided Wilcoxon signed-rank test (followed by Benjamini-Hochberg FDR) was used to compare cell type enrichments in the co-expression submodules in a one-vs-rest fashion. The cell types used in the enrichment analysis were: neurons, TAs, cyclings TAs, myocytes, goblet cells, colonocytes, fibroblasts, glia, lymphatic cells, macrophages, enteroendocrine cells, mesothelial cells, stem cells, T cells, tuft cells, B cells and vascular cells. The maximum adjusted p-value from all comparisons was reported and considered significant if less than 0.05.

### KEGG pathway enrichment

KEGG database[96] gene sets were tested for enrichment in each cell-type specific submodule with a one-tailed Fisher’s exact test followed by a Benjamini-Hochberg FDR. KEGG pathways with FDR <0.05 were visualized.

## Code availability

All code is deposited on GitHub (https://github.com/brittalot/spatial_host_microbiome_sequencing).

## Acknowledgements

We thank Ania Hupalowska for help with figure preparation. We thank Eric Brown for help with mouse husbandry and Hera Vlamakis for fruitful discussions. Work was supported by the Knut and Alice Wallenberg Foundation, Beijer Laboratory for Gene and Neuro Research, the Royal Swedish Academy of Sciences and Swedish Society for Medical Research (S.V.), the Hans Werthén Foundation, Foundation Blanceflor Boncompagni Ludovisi, née Bildt (B.L), the Klarman Cell Observatory, the Manton Foundation, and HHMI (A.R.). S.V was supported as a Wallenberg

Fellow at the Broad Institute of MIT and Harvard and as a Wallenberg Academy Fellow at Uppsala University. A.R. was an Investigator of the Howard Hughes Medical Institute.

## Author contributions

S.V. conceived and designed the study and experiments with guidance from A.R.; B.L. performed the experiments; S.V. and B.L. analyzed the data; B.L. built the deep learning model with help from M.S.; S.V. annotated the histological sections. S.V., B.L. and A.R. wrote the manuscript with input from all the authors. All authors discussed the results.

