## Supplementary information for "Spatial host-microbiome sequencing"

### Supplementary Figures

Supplementary Figure 1

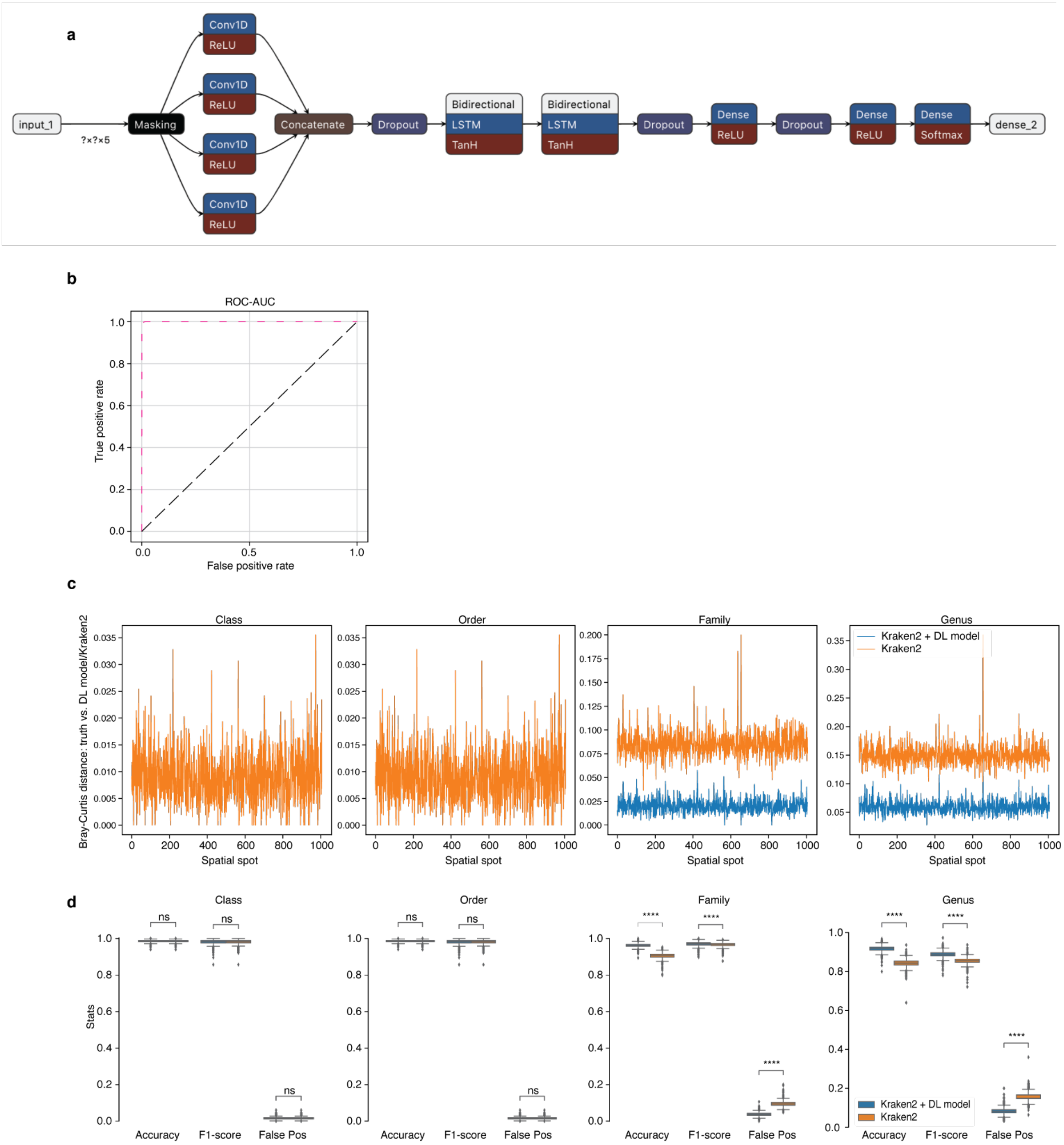


**Supplementary Fig1. Deep learning model performance. (a)** Overview of the DL model architecture. The model uses five-dimensional binary vectors to represent the nucleotides and these are first masked to ignore padded entries, followed by four layers of a one-dimensional convolutional layer with increasing kernel-sizes. After the convolutional layers, a concatenation and a dropout followed. The model then used two bidirectional Long Short Term Memory networks to process the sequences in both directions, before another dropout layer. This was followed by first a dense layer (reLU activation), second a dropout layer, third another dense layer (reLU activation) and finally a dense layer (softmax activation). **(b)** ROC curve illustrating the performance of the DL model on 20% of the test data. **(c)** Bray-Curtis dissimilarities between true data labels and the same data taxonomically classified using either Kraken2 alone (orange) or Kraken2 + DL model (blue) per spatial spot on four taxonomic levels. **(d)** Accuracy, F1 score and false positive rate for simulated sequencing data classified using either Kraken2 alone (orange) or Kraken2 + DL model (blue) on four taxonomic levels (n = 3). Statistical significance markings (two-sided *t*-test) are displayed; $0.05<p\leq1$ (ns), $p\leq0.0001$ (****).

Supplementary Figure 2

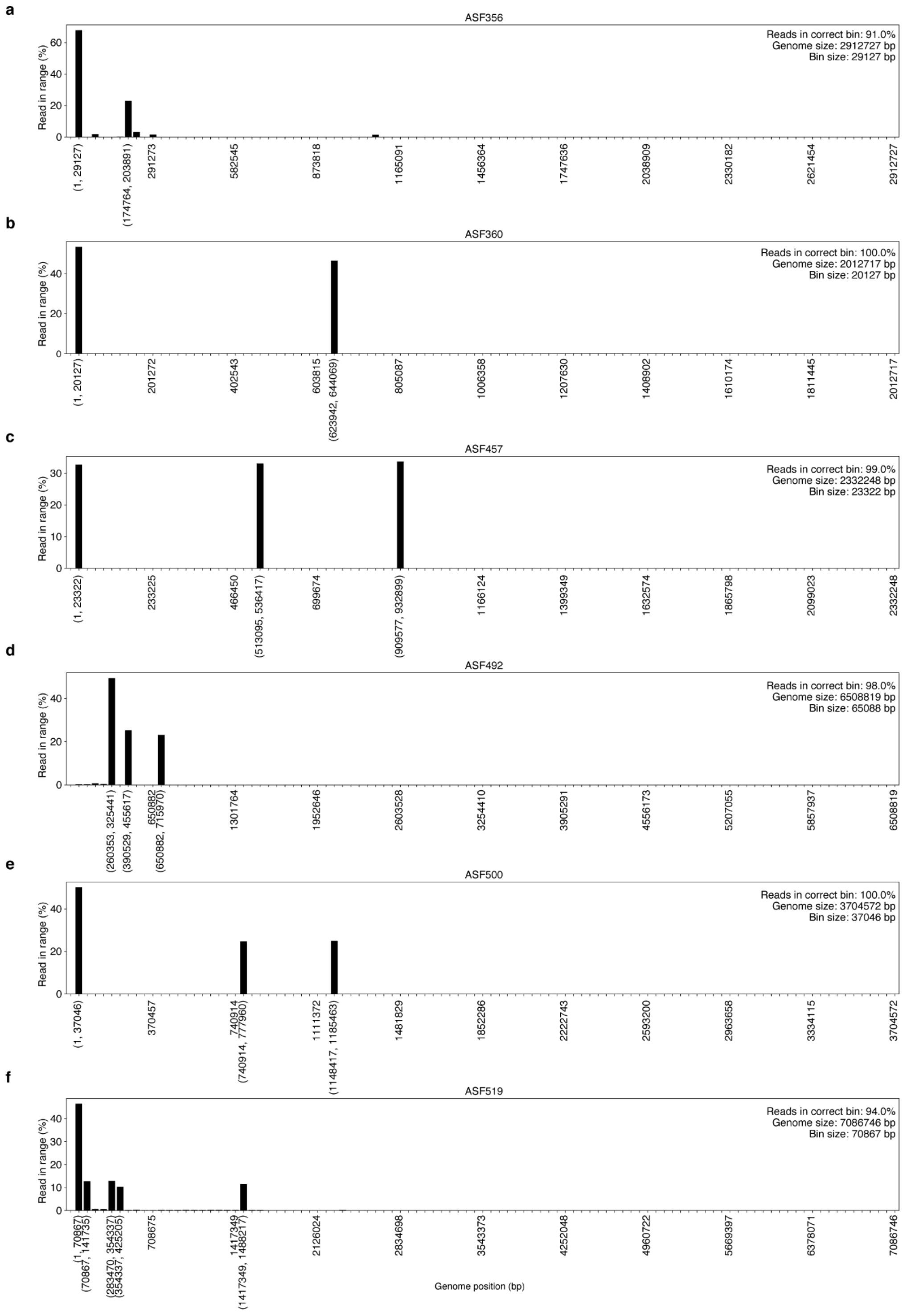


**Supplementary Fig2. Reads captured by the 16S surface probe align to the expected 16S rRNA gene locations in selected bacteria in the ASF mouse model.** Genome locations of proportions of binned read alignments in **(a)** ASF365, **(b)** ASF360, **(c)** ASF457, **(d)** ASF492, **(e)** ASF500 and **(f)** ASF519. The x axis represents genomic positions of the bins and the start and end position of each bin where the proportion of detected reads was over 10% is shown in brackets.

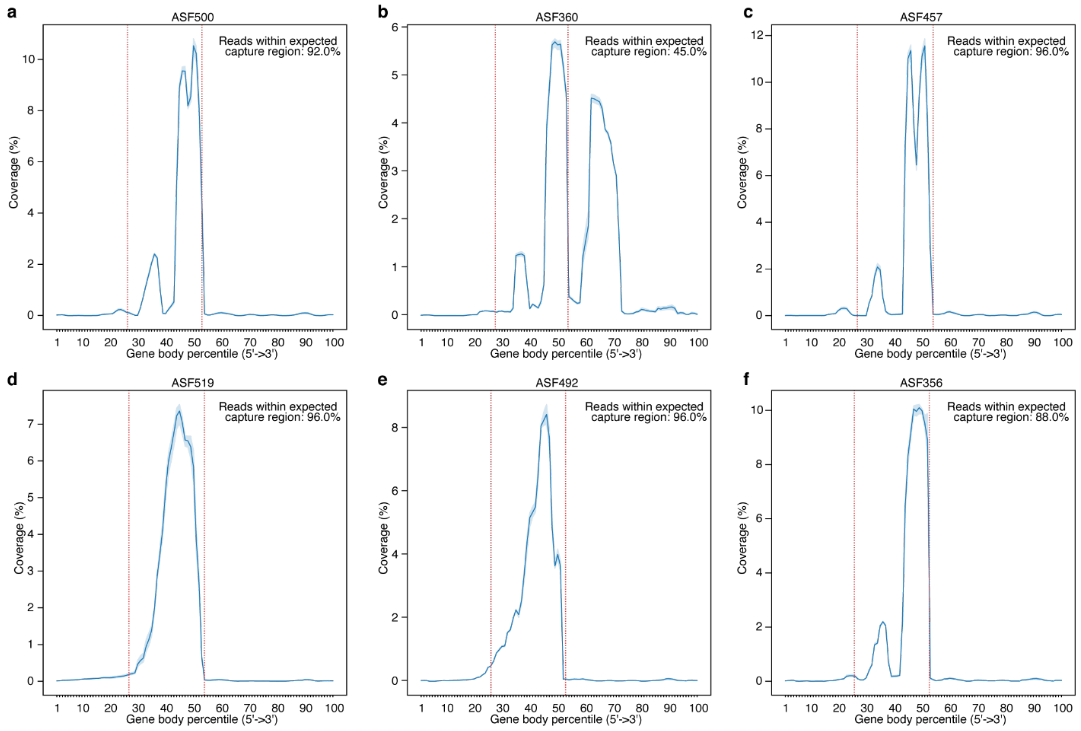


**Supplementary Fig3. 16S surface probe captured the expected part of the 16S rRNA gene in selected bacteria in the ASF mouse model.** Mean gene body coverage (n = 3) over 16S rRNA genes with 95% confidence interval shown in **(a)** ASF500, **(b)** ASF360, **(c)** ASF457, **(d)** ASF519, **(e)** ASF492 and **(f)** ASF356. Red vertical lines mark the expected captured region of the 16S rRNA gene.

Supplementary Figure 4

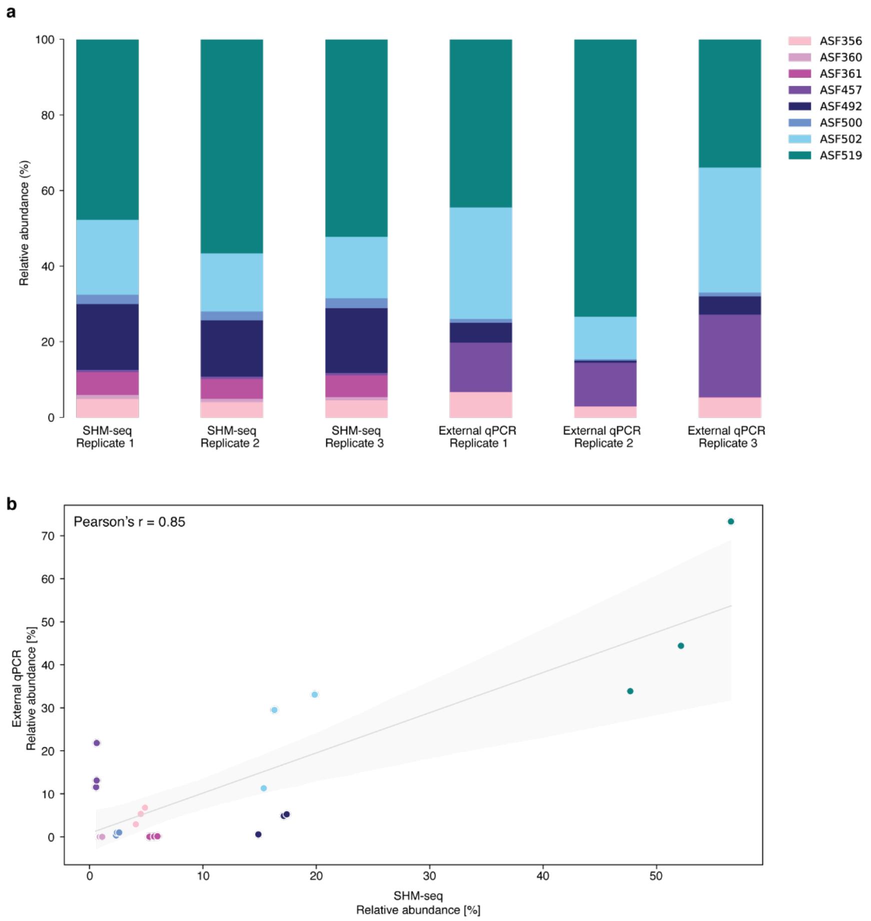


**Supplementary Fig4. Bacterial alignment rates in ASF mice obtained using SHM-seq. (a)** Relative abundances of bacterial reads from ASF tissue sections (n = 3) aligned to the ASF mouse genome reference (SHM-seq) in comparison with reference RT-qPCR count data[1] (external reference). **(b)** Comparison of bacterial read abundances from ASF tissue sections (n = 3) aligned to the ASF mouse genome reference (x axis) with reference RT-qPCR count data[1] (y axis). Color coding is shared between the panels. Shaded areas: 95% confidence interval.

Supplementary Figure 5

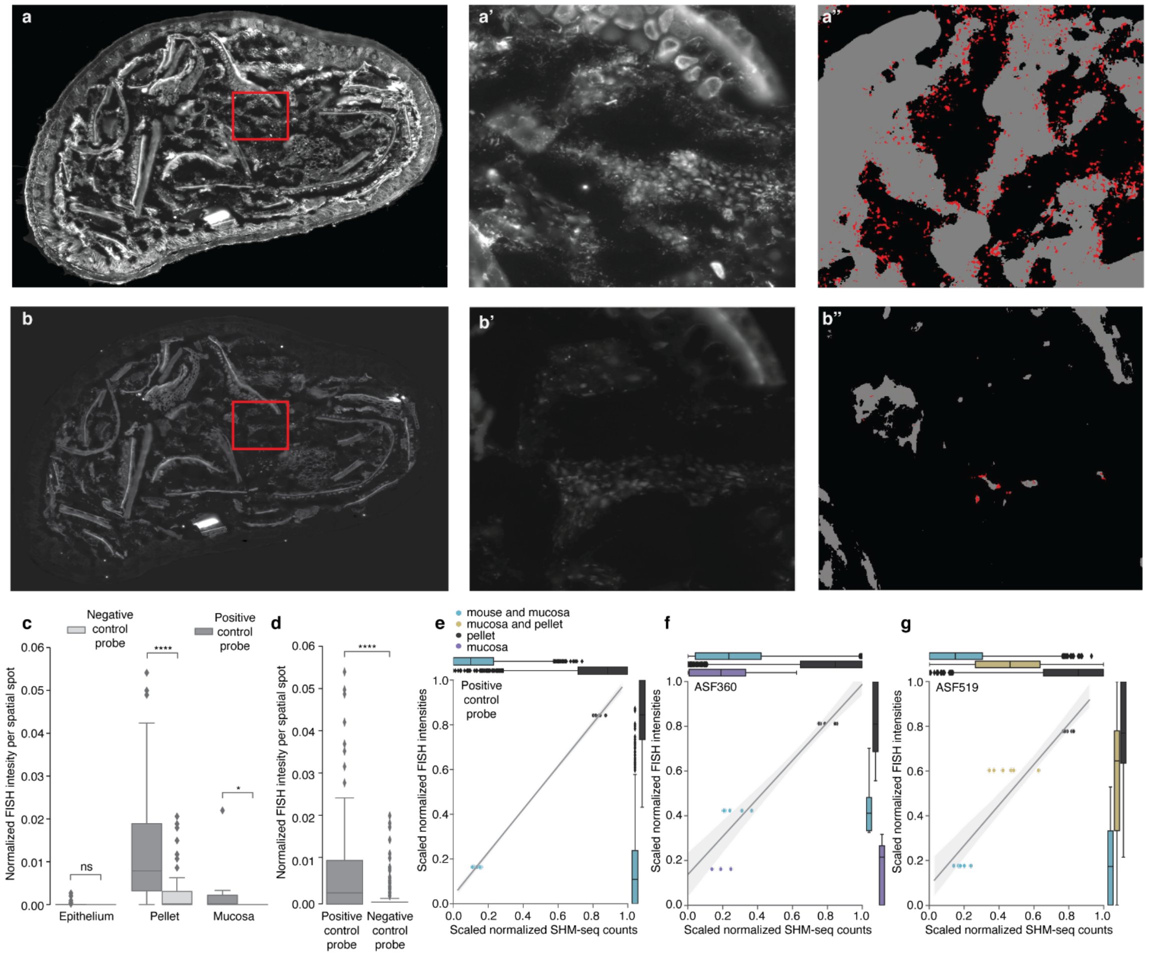


**Supplementary Fig5. Fluorescence *in situ* hybridization of select bacterial targets in ASF colon sections. (a)** and **(a')** Raw fluorescent signal from a FISH experiment using a positive control probe; cross section of an ASF mouse colon and zoomed-in region, respectively. **(a'')** Same area as in **(a')** but with bacterial locations noted in red and fibers noted in gray as classified using Ilastik processing (**Methods**). **(b)** and **(b')** Raw fluorescent signal from a FISH experiment using the negative control probe; cross section of an ASF mouse colon and zoomed-in region, respectively. **(b'')** Same area as in **(b')** but with bacterial locations noted in red and fibers noted in gray as classified using Ilastik processing (**Methods**). **(c)** Box plot of normalized bacterial fluorescence intensity per spatial spot in three MROIs for the positive (dark gray) and negative (light gray) control probes. **(d)** Box plot of normalized bacterial fluorescence intensities per spatial spot for the positive (left) and negative (right) control probes. **(e, f, g)** Box plot and scatter plot with linear regression model fit of scaled normalized fluorescence intensity from FISH experiment (y axis) *vs*. scaled normalized bacterial count generated using SHM-seq (x axis) in shared MROIs **(Methods)** using **(e)** positive control FISH probe, **(f)** FISH probe targeting ASF360 and **(g)** FISH probe targeting ASF519. **(e, f, g)** Boxplots show the spread of normalized signals per region. Scatterplot shows the mean signal per region and sample (n = 6). Shaded areas: 95% confidence interval. Legend in **(e)** is shared between subfigures. **(c, d)** Statistical significance markings (two-sided *t*-test) are displayed; $0.05<p\leq1$ (ns), $0.01<p\leq0.05$ (*), $p\leq0.0001$ (****).

Supplementary Figure 6

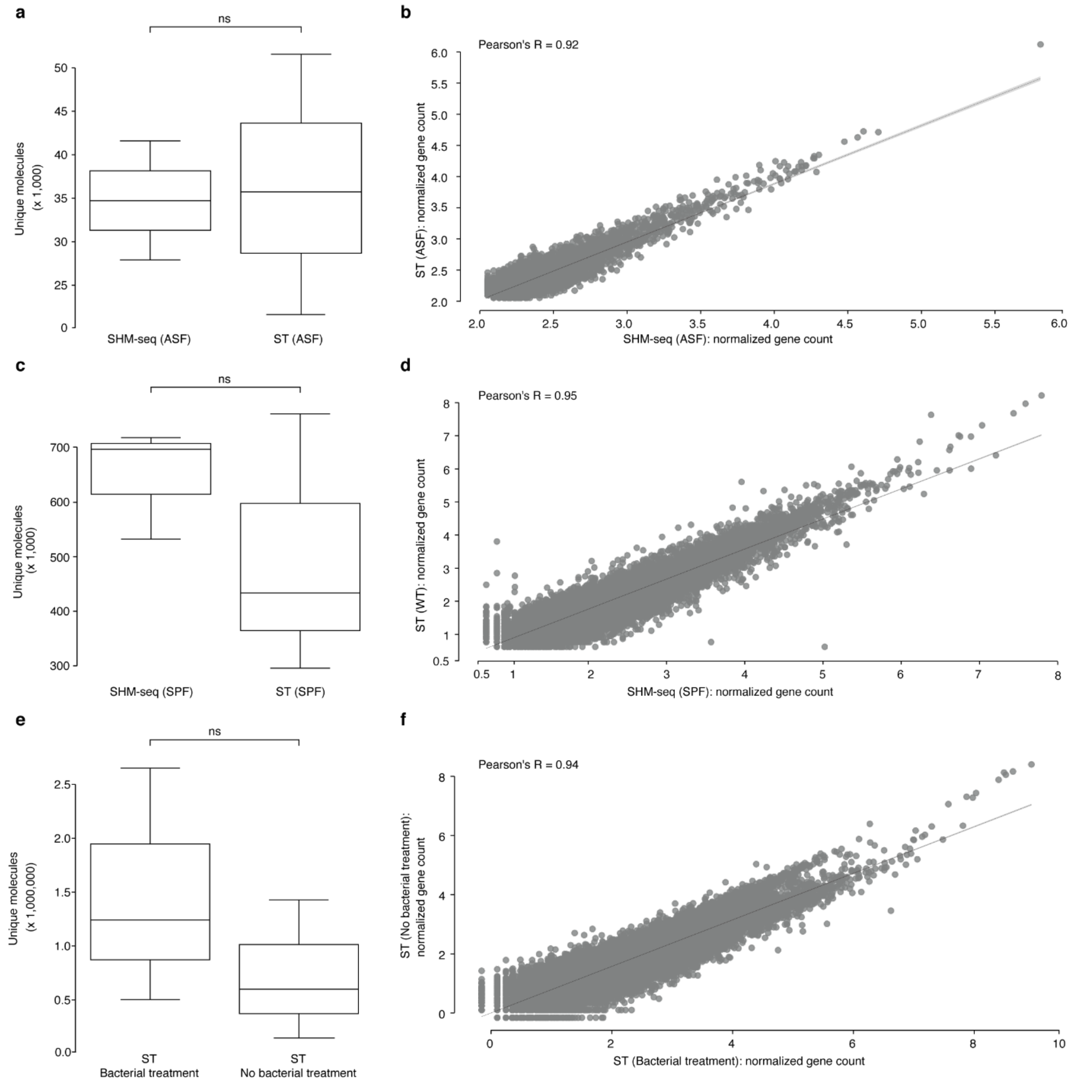


**Supplementary Fig6. Overall spatial gene expression performance metrics in SPF and ASF mouse tissue sections. (a)** Comparison of the total number of UMIs obtained after sequencing in the ASF mouse model using SHM-seq (left, n = 3) *vs.* ST (right, n = 3). **(b)** Correlation between the normalized gene expression counts in the ASF mouse model using SHM-seq (x axis, n = 3) and ST (y axis, n = 3). **(c)** Comparison of the total number of UMIs obtained after sequencing in the SPF mouse model using SHM-seq (left, n = 3) *vs.* ST (right, n = 3). **(d)** Correlation between the normalized gene expression counts in the SPF mouse model using SHM-seq (x axis, n = 3) and ST (y axis, n = 3). **(e)** Comparison of the total number of

UMIs in the SPF mouse model obtained after sequencing with (left green, n = 3) and without (right, n = 3) applying bacterial treatment in ST experiments **(Methods)** *in situ*. **(f)** Correlation between normalized gene expression in the SPF mouse model with (x axis, n = 3) and without (y axis, n = 3) applying bacterial treatment in ST experiments **(Methods)** *in situ*. Statistical significance markings (two-sided *t*-test) are displayed; $0.05<p\leq1$ (ns).

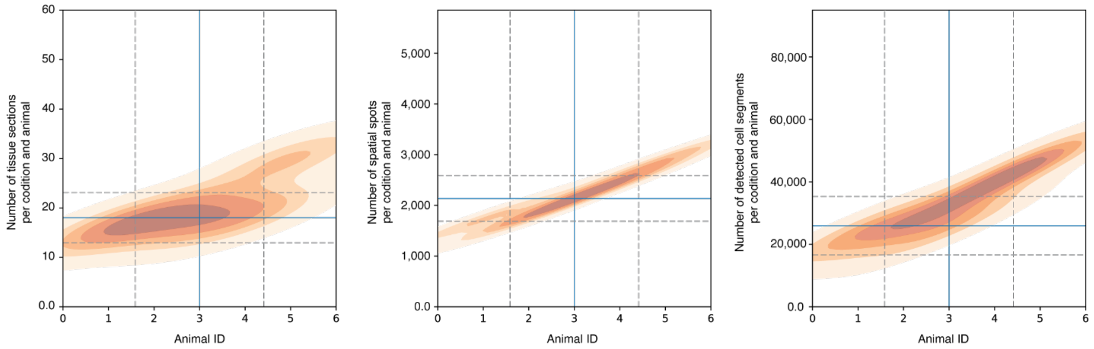


**Supplementary Fig7. Sampling metrics for 100 colonic mouse sections (SPF and GF mice). (a)** Density scatter plots showing the number of tissues sampled in each mouse model and animal (y axis, left), number of spatial spots sampled in each mouse model and animal (y axis, middle), and number of cell segments detected in each mouse model and animal (y axis, right) *vs.* number of animals sampled per condition (x axes). Blue lines; median number of observations per condition, gray dashed lines; standard deviations for all observations per condition.

### Supplementary Figure 8

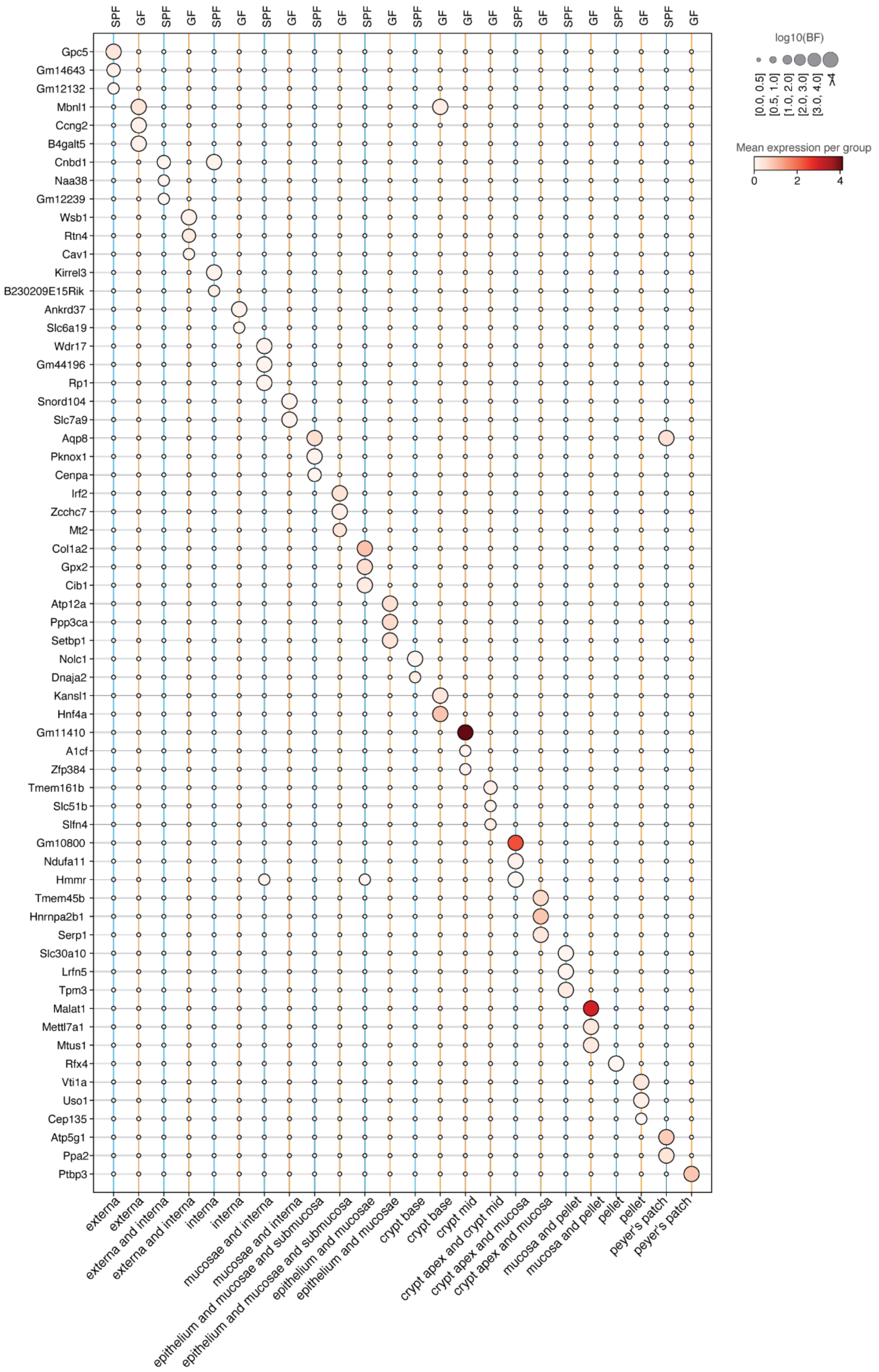


**Supplementary Fig8. Spatial gene expression difference per mouse model and MROI.** Top 3 differentially expressed genes (rows) between conditions (GF *vs*. SPF) for each MROI (columns). Spot size corresponds to log10 Bayes Factor (BF) and color scale to the normalized gene expression.

Supplementary Figure 9

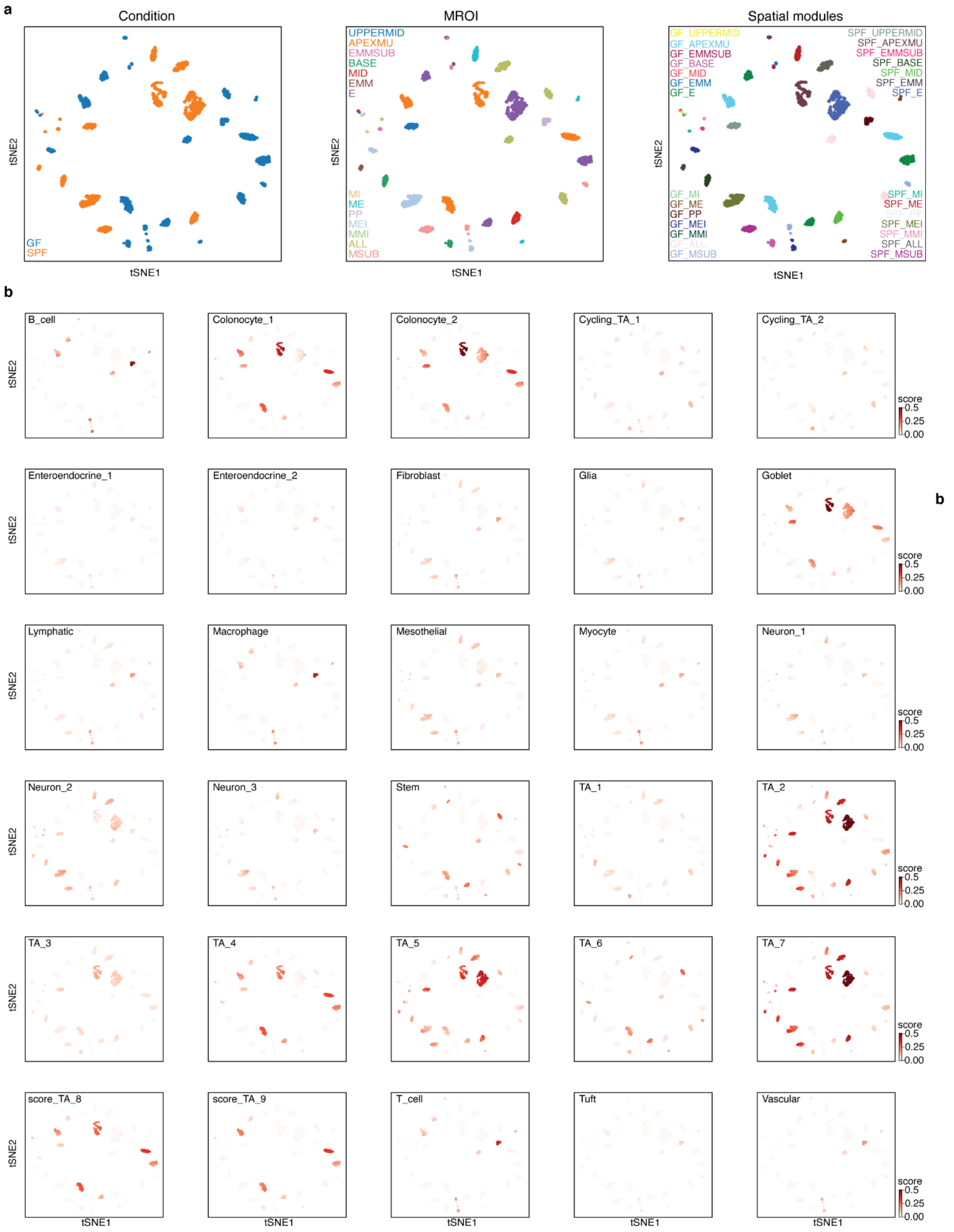


**Supplementary Fig9. t-SNE representations of spatial modules. (a)** t-SNE representations of mouse conditions (left), MROIs (middle) and spatial modules (right). **(b)** t-SNE representations of constituent cell types (color scale) present in each of the spatial modules.

**Supplementary Tables**

**Supplementary Table 1. Cell-type specific submodule expression.**

**Supplementary Table 2. KEGG pathway analysis for all modules and submodules.**

### SUPPLEMENTARY INFORMATION REFERENCES

1. Sarma-Rupavtarm, R. B., Ge, Z., Schauer, D. B., Fox, J. G. & Polz, M. F. Spatial distribution and stability of the eight microbial species of the altered schaedler flora in the mouse gastrointestinal tract. *Appl. Environ. Microbiol.* **70**, 2791–2800 (2004).
